# Dysregulated interactions triggered by a neuropathy-causing mutation in the IPV motif of HSP27

**DOI:** 10.1101/708180

**Authors:** T. Reid Alderson, Elias Adriaenssens, Bob Asselbergh, Iva Pritišanac, Heidi Y. Gastall, Marielle A. Wälti, John M. Louis, Vincent Timmerman, Andrew J. Baldwin, Justin L. P. Benesch

## Abstract

HSP27 (HSPB1) is a systemically expressed human small heat-shock protein that forms large, dynamic oligomers and functions in various aspects of cellular homeostasis. Mutations in HSP27 cause Charcot-Marie-Tooth disease, the most common inherited disorder of the peripheral nervous system. A particularly severe form of the disease is triggered by the P182L mutation within the highly conserved IxI/V motif of HSP27. Here, we observed that the P182L variant of HSP27 lacks the ability to prevent the aggregation of client proteins and formed significantly larger oligomers both *in vitro* and *in vivo*. NMR spectroscopy revealed that the P182L IxI/V motif binds its α-crystallin domain with significantly lower association rate, and thus affinity, rendering the binding site more available for other interactors. We identified 22 IxI/V-containing proteins that are known to interact with HSP27 and could therefore bind with enhanced affinity to the P182L variant. We validated this hypothesis through co-immunoprecipitation experiments, revealing that the IxI/V motif-bearing co-chaperone BAG3 indeed binds with higher affinity to the P182L variant. Our results provide a mechanistic basis for the impact of the P182L mutation on HSP27, and highlight the general importance of the IxI/V motif and its role in protein-protein interaction networks.

## Introduction

HSP27 (HSPB1) is a systemically expressed human small heat-shock protein (sHSP) that performs diverse functions under basal and stressful cellular conditions (1–3). With significant roles in the maintenance of protein homeostasis, regulation of the redox environment, prevention of apoptosis, and stabilization of the cytoskeletal network, the biological activity of HSP27 is critical to overall cellular health (4–8). Dysregulation of the activity or expression of HSP27 can result in debilitating diseases, including cancers (9, 10), neurodegenerative diseases (11), and neuropathies (12). Over 30 heritable HSP27 mutations are implicated in Charcot-Marie-Tooth (CMT) disease (13–15), a group of neuropathies that affects ca. 1 in 2500 individuals and is the most common inherited disorder involving the peripheral nervous system (16, 17). CMT disease is characterized by progressive demyelination (type 1 CMT), axonal loss (type 2 CMT), or their combination (18). When the affected axons exclusively include motor neurons, the disease is referred to as distal hereditary motor neuropathy (dHMN) (18). Transgenic HSP27 mouse models develop CMT disease-like symptoms (19), thereby implicating HSP27 as a direct driver of CMT disease onset.

HSP27 contains three domains: a conserved α-crystallin domain (ACD) that is flanked by an N-terminal domain (NTD) and C-terminal region (CTR), which is disordered and contains a highly conserved IxI/V motif. Intermolecular contacts between all three domains (4, 20) facilitate the assembly of HSP27 into large, dynamic oligomers with an average mass near 500 kDa (21–25). In mammalian sHSPs, the central residue in the IxI/V motif is generally proline and this short linear motif (SLiM) facilitates binding to the structured ACD: high-resolution structures have been obtained for the isolated ACD (22, 26) bound to a peptide containing the IxI/V motif (22). SLiMs ranging from 2-12 residues in length can mediate interactions between disordered regions and structured domains (27, 28), and disease-causing mutations within disordered regions often impact SLiMs or create new ones (27, 29). The IxI/V SLiM within the disordered CTR of sHSPs reversibly transitions between a flexible, unbound form and a rigid, ACD-bound state (25). Despite mediating the ACD-CTR interaction, the IxI/V motif is not necessarily a prerequisite for oligomeric assembly in HSP27 or the related sHSPs αA- and αB-crystallin. Indeed, mutations that attenuate or prevent IxI/V binding did not decrease the average oligomeric size (30–35). In addition, the IxI/V motif can facilitate the interaction of HSP27 with other proteins that contain this motif, including the HSP70 cochaperone Bcl-2-associated anthanogene-3 (BAG3), which plays a significant role in regulating apoptosis and autophagy (36). BAG3 contains two IPV motifs, and the IxI/V-mediated BAG3-HSP27 interaction leads to formation of an HSP27-BAG3-HSP70 ternary complex, thus providing a physical link between HSP27 and HSP70 that allows client transfer to occur (37).

The majority of the CMT disease-causing mutations in *HSPB1* reside in the ACD (15, 38), with fewer sites located in the NTD (39) and CTR (40). Notably, two CMT-causing mutations lie within the highly conserved IxI/V motif (P182L, P182S) with additional mutations located nearby (12, 15). The P182L variant comprises one of the most clinically severe HSP27 variants, with symptoms manifesting in the first five years of life, whereas mutations in the ACD generally result in adult-onset symptoms (41). However, the molecular basis of the disease-causing mutations involving the P182 codon in HSP27 remains unknown. Within the IxI/V motif, P182 is the only residue that is implicated in CMT disease; other mutational sites in the ACD besides a frameshift at S158 do not reside in the β4 or β8 strands that comprise the IxI/V binding site.

Here, we sought to understand the biophysical significance of the P182L mutation in HSP27 and its impact on the interaction of the CTR with the ACD. We observed that the P182L mutation significantly increased the average molecular mass of soluble HSP27 oligomers and disrupted its ability to prevent substrate aggregation. The binding of the IxI/V motif to the ACD was studied in detail with NMR spectroscopy, which revealed that the affinity of the P182L variant is decreased by nearly one order of magnitude due to a lower association rate. A consequence of the weakened affinity for the ACD is a more exposed binding site for other IxI/V-containing HSP27-interacting proteins. We find that BAG3, an HSP70 co-chaperone with two IPV motifs, indeed binds with enhanced affinity for the P182L variant of HSP27. Given the highly conserved nature of the IxI/V motif, our findings should be generally relevant for sHSPs that harbor this tripeptide motif.

## Results

While the majority of CMT-associated mutations in HSP27 reside in the structured ACD (Fig. 1A), mutations within the disordered NTD and CTR also cause CMT disease. One of the most severe phenotypes is associated with the P182L variant, which falls in the intrinsically disordered CTR and changes the highly conserved Pro residue within the IxI/V motif (Ile-Pro-Val) to a Leu (Fig. 1B). The pathogenicity caused by mutation of this SLiM has been demonstrated in a transgenic mouse model (19), but the biophysical and mechanistic basis of the P182L mutation remain unclear. To shed light on the structural impact of the P182L mutation (Fig. 1A, Fig. 1B), we investigated the biophysical and functional properties of the wild-type (WT) and P182L variants of full-length HSP27. Our experiments also utilize the isolated ACD (Fig. 1C) to characterize the binding of the IxI/V motif in its WT and P182L forms. The ACD folds into a β-strand-rich dimer (Fig. 1D) and the IxI/V motif binds to the ACD in the groove located between the β4 and β8 strands (Fig. 1D) through hydrophobic interactions and hydrogen bonds to the backbone atoms of V111, T113, and L157 (Fig. 1E). In HSP27, the IxI/V motif corresponds to I181-P182-V183; the Ile and Val residues penetrate the β4/β8 groove while P182 does not make any direct contacts with the ACD (Fig. 1E).

**Fig. 1.**
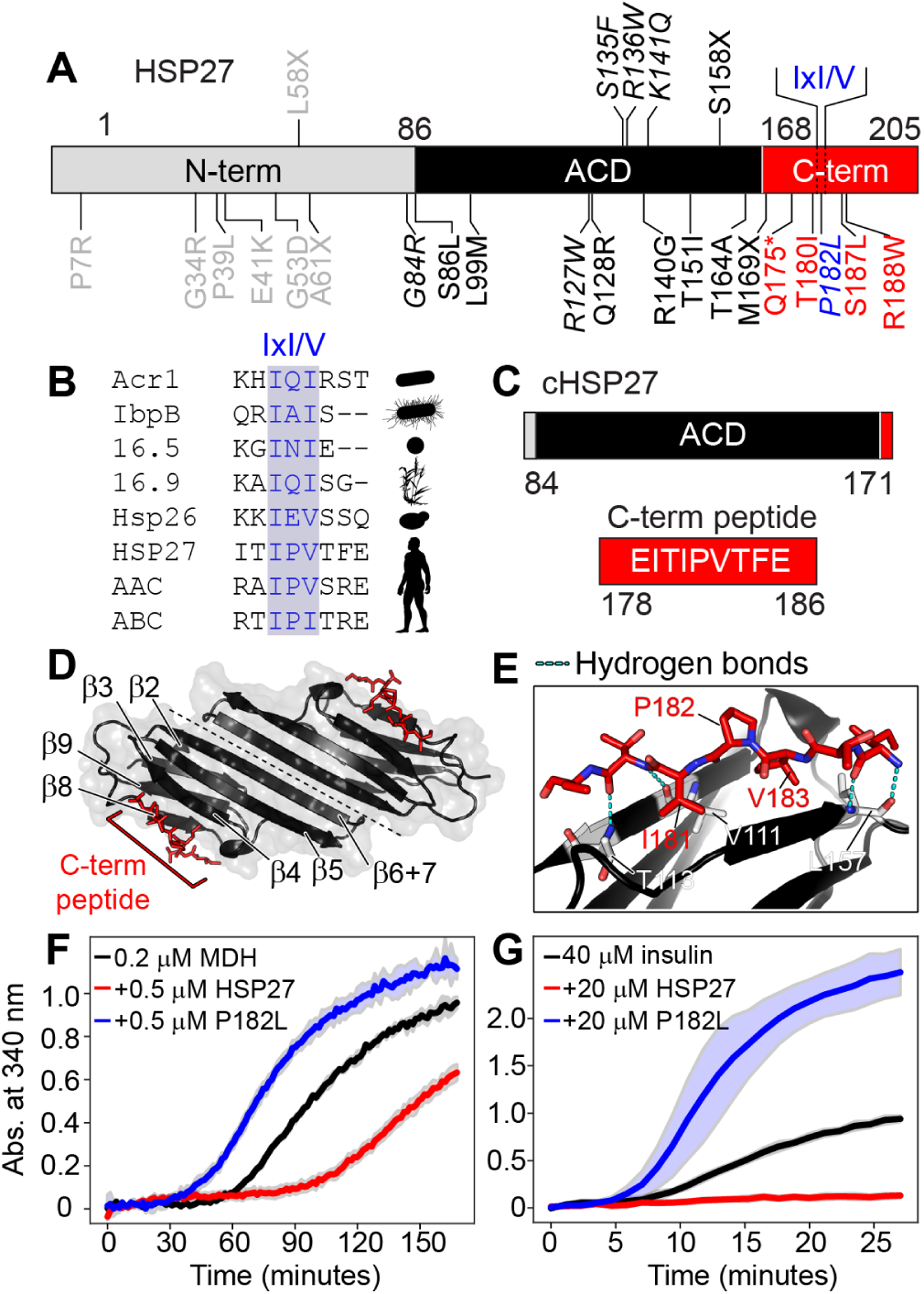
The P182L mutation is located in the IxI/V motif of HSP27 and disrupts its chaperone activity. (**A**) Domain architecture of human HSP27: N-terminal domain (N-term), α-crystallin domain (ACD), and C-terminal region (C-term). The conserved IxI/V motif in the C-terminal region is indicated. Missense, nonsense, and frameshift mutations associated with CMT disease and dHMN are shown. The P182L mutation in the IxI/V motif is blue. Residues with multiple mutations are italicized (e.g. P182L, P182S). The asterisk denotes a nonsense mutation and the X indicates a frameshift mutation. (**B**) Sequence alignment of IxI/V motifs and adjacent residues from the sHSPs Acrl (*M. tuberculosis*), IbpB (*E. coli*), Hsp16.5 (*M. janaschii*), Hsp16.9 (*T. aestivum*), Hsp26 (*S. cerevisiae*), and human HSP27, αA-crystallin, and αB-crystallin. (**C**) Domain boundaries of the ACD used in this study (cHSP27) and residues in the C-terminal peptide that contains the IxI/V motif. (**D**) Three-dimensional structure of the ACD dimer (black) bound to a peptide from the C-terminal region (red) containing the IxI/V motif. The β-strands of the ACD are numbered. (**E**) Zoomed-in region from D showing the contacts made between the C-terminal peptide and the ACD. (**F** MDH (0.2 µM) was incubated at 40 °C in the absence (black) or presence of 0.5 µM WT HSP27 (red) or 0.5 µM P182L HSP27 (blue). (**G**) Insulin (40 µM) was incubated at 40 °C in the absence (black) or presence of 20 µM WT HSP27 (red) or 20 µM P182L HSP27 (blue). The y-axes depict the absorbance at 340 nm due to the formation of large aggregates. The solid lines represent the average of three replicates with the filled area reflecting ± one standard deviation.

### The P182L variant has impaired chaperone activity *in vitro*

We purified recombinant human HSP27 and the CMT-implicated P182L variant from *E. coli*, and compared their chaperone activities *in vitro* using a substrate aggregation assay. We tested their ability to prevent the aggregation of two model substrate proteins, malate dehydrogenase (MDH) and insulin. The WT form of HSP27 proved highly effective at inhibiting the aggregation of both substrates (Fig. 1F, Fig. 1G, red traces), indicative of its potent anti-aggregation activity. However, the P182L variant was unable to prevent the aggregation of either substrate (Fig. 1F, Fig. 1G, blue traces), revealing a drastic loss of function *in vitro*. Indeed, the P182L form of HSP27 appeared to co-aggregate with MDH and insulin, based on the elevated light scattering in the presence of the chaperone. Similar co-aggregation was previously observed for the cardiomyopathy-causing R120G variant of αB-crystallin (42). Control aggregation assays using P182L or HSP27 alone indicated the absence of appreciable aggregation (Supplementary Fig. 1), suggesting that P182L is stable in solution and the loss of P182L chaperone activity is not due to self-aggregation. The P182L mutation thus significantly alters the ability of HSP27 to prevent the aggregation of client proteins.

### P182L forms large oligomers both *in vitro* and *in vivo*

We next characterized the disease-causing P182L variant with size exclusion chromatography coupled to multi-angle light scattering (SEC-MALS) to determine the average mass and hydration radius *in vitro*. For WT HSP27 and its isolated ACD, the SEC-MALS data revealed average masses near 500 and 20 kDa (Fig. 2A), respectively, which are consistent with previous data. In contrast, the SEC-MALS profile of the P182L variant revealed significantly larger oligomers that exhibited an average mass near 14 MDa (Fig. 2A), nearly a 30-fold increase over the WT protein. The average radius of hydration (*R*_*h*_) of the P182L variant increased from *ca*. 9 nm to *ca*. 38 nm. With a monomeric mass of 22.7 kDa, the P182L variant therefore contains an average of *ca*. 620 subunits, whereas the WT protein comprises *ca*. 24 subunits. In support of these SEC-MALS data, negative-stain electron microscopy (EM) demonstrated that the P182L variant assembles into much larger oligomeric forms than the WT protein (Fig. 2B). The polydispersity of oligomers populated by both the P182L variant and the WT protein was also evident from the negative-stain EM images (Fig. 2B). Together, the SEC-MALS and negative-stain EM data establish that the disease-causing P182L mutation drastically shifts the oligomeric distribution of HSP27 toward much larger species.

**Fig. 2.**
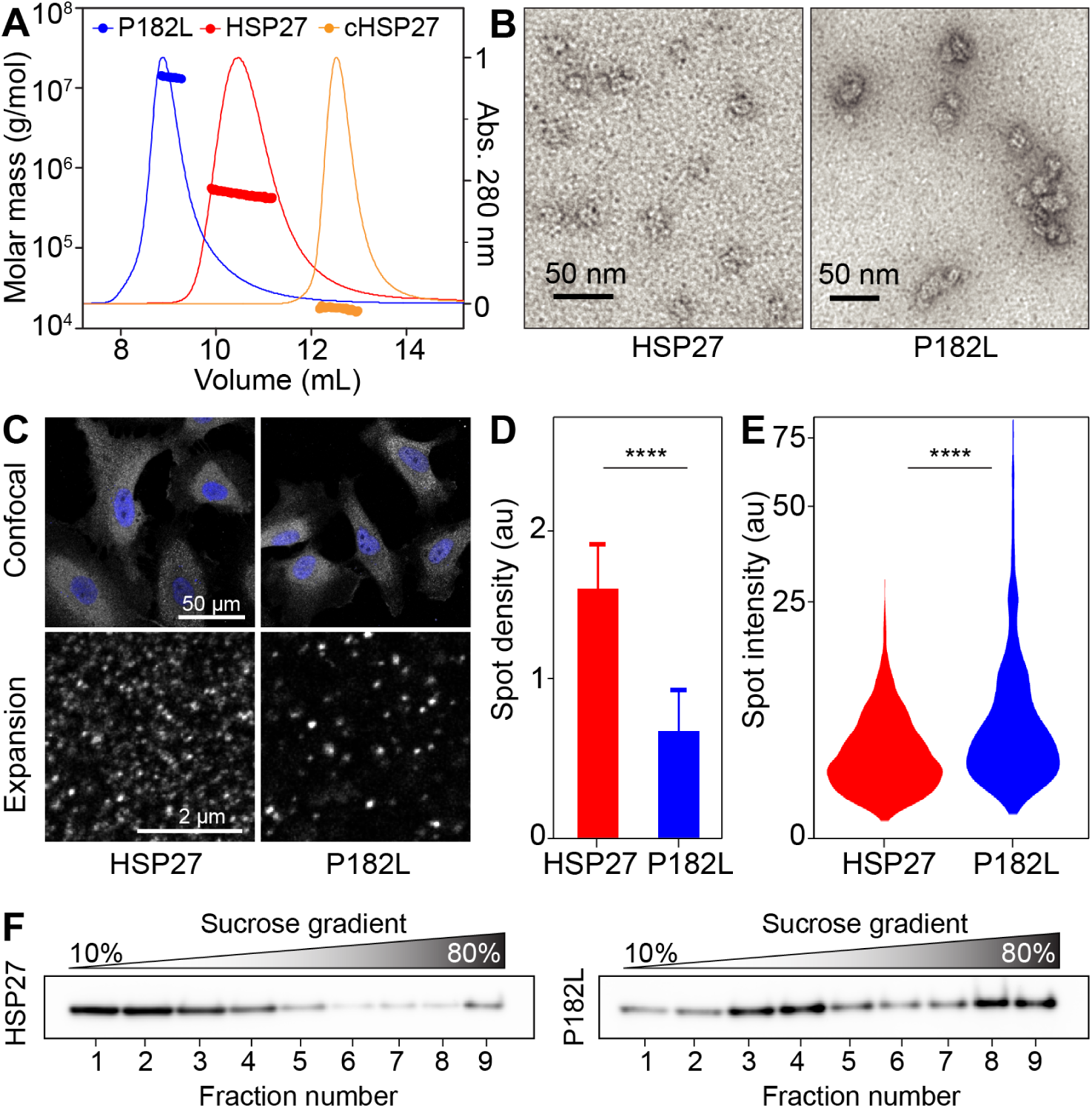
The P182L variant of HSP27 forms larger oligomers both *in vitro* and *in vivo*. (**A**) SEC-MALS data from WT HSP27 and the P182L variant. Both samples were injected at 40 µM concentration (monomer) in 30 mM sodium phosphate, 2 mM EDTA, 100 mM NaCl buffer at pH 7. The UV signals are plotted on the right y-axis as solid lines and the molecular masses are plotted on the left y-axis as circles. (**B**) Negative-stain EM images show that the P182L variant (left) forms much larger oligomeric assemblies than the WT form (right). Both samples were loaded on the grids at the same concentration (4.5 µM monomer concentration). (**C**) Expression of V5-epitope-tagged HSP27 (WT or P182L mutant) in HSP27 knock-out HeLa cells and immunostaining of HSP27. Expansion microscopy increases resolution and allows resolution of the distribution of individual soluble HSP27 assemblies in the cytoplasm. The scale bars correspond to the non-expanded dimensions, derived by dividing by the sample expansion factor of 4.3. (**D**) The P182L mutation decreases the density of HSP27 spots upon detection of local fluorescence intensity maxima and quantification of the number of detected spots in expanded samples (n = 30). The error bar is the standard deviation for the different cytoplasmic regions. The asterisks correspond to a P value < 0.0001 as obtained from a t-test. (**E**) Distribution of the intensity of individual spots (n = 3342) after Gaussian fitting of fluorescence intensity in the neighborhood of detected spots. The asterisks correspond to a P value < 0.0001 as obtained from a Mann-Whitney U test. (**F**) Fractionation of HeLa cells overexpressing either HSP27 or the P182L mutant by a 10% to 80% sucrose gradient. Western blot against anti-V5 is shown.

The above experiments made use of protein samples that were recombinantly expressed and purified from *E. coli*. The oligomeric state of HSP27 and the P182L variant in human cells has also been reported to be altered, as overexpression of the P182L mutant to leads to increased formation of large insoluble HSP27-containing cytoplasmic aggregates (43–45). Typically, higher levels of mutant HSP27 expression resulted in higher percentages of aggregate-containing cells and vice versa, with the size of individual cellular aggregates strongly variable and extending to tens of microns (data not shown). Indeed, upon transient transfection of V5-epitope-tagged P182L HSP27 in HeLa cells, we found that more than 25% of cells contained large, high-intensity aggregates visible by conventional microscopy, whereas comparable transient expression of wild-type HSP27 caused aggregation in less than 2% of cells (Supplementary Fig. 2). To also visualize these soluble oligomeric states of HSP27 *in vivo*, we performed expansion microscopy, which allows imaging of HSP27 at the nanoscale and analysis of the distribution of soluble HSP27 in the cytoplasm. Expansion microscopy increases the effective resolution by more than 4-fold compared to conventional microscopy (46). As a result, individual soluble HSP27 assemblies that are not distinguishable using standard confocal microscopy could be visualized and subsequently analyzed (Fig. 2C). Comparing the number of detectable spots per cytoplasmic volume using expansion microscopy shows that expression of the P182L mutant leads to a drastic reduction in the density of HSP27 assemblies (Fig. 2D). Furthermore, the fluorescence intensity of each spot is determined by the number of HSP27 subunits, and therefore indirectly reflects the size of the oligomeric assemblies. Our measurements indicated that the P182L mutation increases the proportion of spots with higher intensities (Fig. 2E). Combined, these results show that assemblies of the P182L mutant are larger and more sparsely distributed in the cytoplasm, consistent with a higher oligomeric state *in vivo*.

To confirm these results by an independent method, we extracted whole cell protein lysates from HeLa cells stably overexpressing WT HSP27 or the P182L mutant and separated these protein lysates over a sucrose gradient. The respective fractions were loaded on western blot and demonstrated a clear shift towards larger protein complexes for the P182L mutant (Fig. 2F). Interestingly, two distinct pools of enriched large complexes were observed in P182L lysates: both the highest fractions containing the large insoluble aggregates (fractions 8-9) as well as larger assemblies of the smaller soluble complexes of HSP27 (fractions 1-2 from WT to fractions 3-4 for P182L). This is consistent with our observations using light microscopy as Supplementary Fig. 2 showed the enrichment of large insoluble aggregates (reflected by the enrichment in fractions 8-9 from the sucrose gradient) while our expansion microscopy indicated that there is also an increase in size of smaller soluble complexes (reflected by the enrichment in fractions 3-4 from the sucrose gradient). Taken together, these biophysical data and functional assays demonstrate that the P182L mutation increases the size of oligomers both *in vitro* and *in vivo* toward very large states that have aberrant activity in preventing protein aggregation.

### The P182L mutation significantly lowers binding affinity for the IxI/V motif

Our results above indicate that the P182L mutation in the IxI/V motif dysregulates oligomeric assembly and chaperone function. To understand the molecular basis of these results, we sought to characterize the binding interaction between the ACD and the IxI/V motifs of the WT form (Ile-Pro-Val) and the P182L variant (Ile-Leu-Val). Previous solution-state NMR studies of WT HSP27 only observed resonances from the disordered CTR, and thus could not probe CTR binding to the ACD in context of the full-length protein (47, 48). We therefore turned to the isolated ACD, which forms stable dimers (2 x 10 kDa) that are known to bind to a peptide encompassing the IxI/V motif of the CTR (20, 22, 31, 33, 34, 37).

With solution-state NMR spectroscopy, we determined the *K*_*d*_ of the ACD binding to a peptide comprising the WT IPV motif (Fig. 3A-C). Since the ACD was uniformly ^15^N labeled, each amide bond in the protein contributed a signal in the 2D ^1^H-^15^N heteronuclear single quantum coherence (HSQC) spectrum (Fig. 3A, B). During the peptide titration, increasing the amount of added peptide led to the disappearance of resonances from ACD residues in the β4/β8 groove (Fig. 3A) and progressive changes in the chemical shifts of other residues (Fig. 3B), i.e. chemical shift perturbations (CSPs). The CSPs (equation 1, Methods) from resonances were globally fit to equation 2 (Methods) to determine the *K*_*d*_ of this binding interaction, yielding a value of ca. 125 *µ*M, which corresponds to a free energy of binding ΔG of −5.3 kcal mol^−1^ at 25 °C (Fig. 3C). Similar results were obtained with the software TITAN (Supplementary Fig. 3) (49). We next measured the binding affinity of the peptide bearing the P182L mutation that is implicated in the onset of CMT (Fig. 3A-C). Surprisingly, even though the central Pro residue in the IPV motif does not directly contact the ACD, its mutation to Leu (P182L) lowers the binding affinity for the ACD by nearly one order of magnitude from *ca*. 125 *µ*M to 1 mM (Fig. 3C), corresponding to a ΔΔG of 1.2 kcal mol^−1^ (from −5.3 kcal mol^−1^ to −4.1 kcal mol^−1^).

**Fig. 3.**
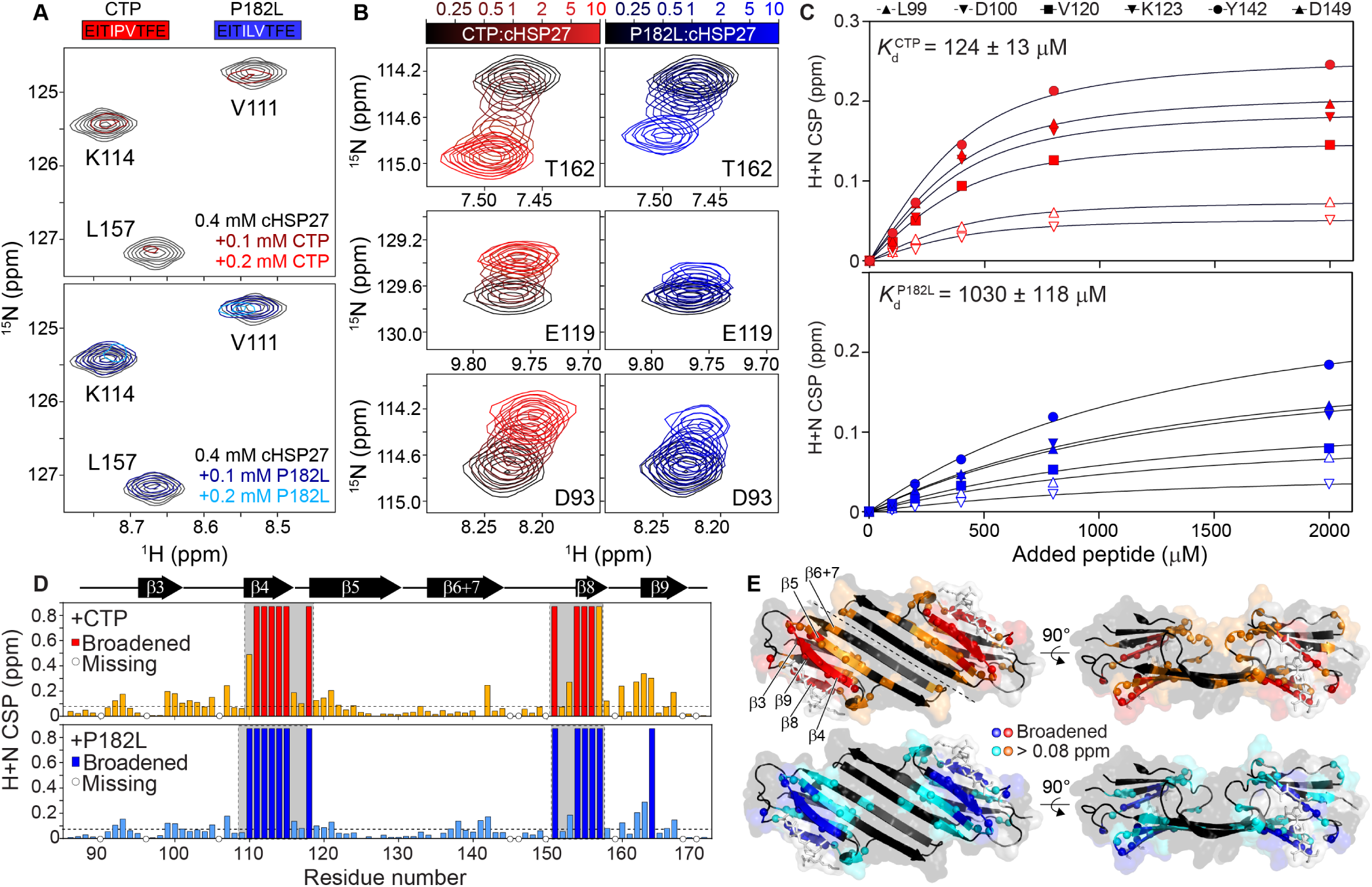
The P182L mutation diminishes the affinity of the IxI/V motif for cHSP27.. (**A**) Primary sequences of the CTP (red) and P182L peptide (blue). The IxI/V motif is indicated with white text. Below: regions of 2D ^1^ H-^15^ N HSQC spectra of ^15^ N-labeled cHSP27 (black) in the presence of increasing amounts of CTP (red) or P182L peptide (blue) at 25 °C. Resonances in the β4 and β8 strands broaden and disappear upon peptide binding, indicative of intermediate exchange. (**B**) Zoomed-in regions of resonances that show fast exchange behavior during peptide binding. The color bars indicate the amount of added WT or P182L peptide. (**C**) The combined ^1^ H^*N*^ and ^15^ N CSPs (eqn 1), are plotted as a function of added peptide concentration. Lines indicate the best fit to eqn 2. The global K_*d*_ +/- one standard deviation is shown in the upper left. (**D**) CSPs shown as a function of residue number for the WT (red) and P182L peptides (blue). (**E**) X-ray structure of cHSP27 bound to its CTP (PDB: 4mjh), shown in white sticks, with the results from (D) plotted onto the structure.

In addition to providing information about the strength of the binding interaction between the IxI/V motif-bearing peptide and the ACD, the NMR data also indicated residues chemical environments changed upon peptide binding. As expected from the crystal structure of the ACD-peptide complex (22), resonances from residues near the β4/β8 groove are significantly impacted by IxI/V binding, becoming severely broadened due to intermediate exchange. In addition, residues in the β5 and β6+7 strands also displayed CSPs (Fig. 3D), which could be caused by subtle structural rearrangements in this region upon peptide binding. In particular, Y142, which makes intermolecular contacts that stabilize the dimer interface, exhibited a CSP that was significantly larger than average. Given the distance of Y142 from the β4/β8 groove, the observation of CSPs for this resonance is suggestive of potential allosteric communication between the β4/β8 groove and the β6+7 strand at the dimer interface (Fig. 3D). A comparison of CSPs between the WT and mutant peptides revealed that many of the same residues were impacted by peptide binding, with residues in the β4/β8 groove also becoming broadened in the P182L mutant (Fig. 3D). The overall similarity of the CSPs between the two peptides suggests that the P182L peptide binds in a similar site as the WT peptide (Fig. 3D), with common allosteric changes to Y142 that manifest upon binding. Therefore, the main difference between the WT and P182L mutant peptides corresponds to a drastically lowered binding affinity of the mutant.

### The P182L IxI/V motif binds with an attenuated association rate

To determine the origin of the weakened binding interaction in the P182L variant, we turned to Carr-Purcell-Meiboom-Gill (CPMG) relaxation dispersion (RD), which enables quantification of the binding kinetics and thermodynamics of protein-ligand interactions (50–54). CPMG RD experiments can determine the kinetics (association and dissociation rates; *k*_*off*_, *k*_*on*_), thermodynamics (population of the bound state; *p*_*B*_), and structural changes (differences in chemical shifts; |Δ*ω*|) involved in the interconversion between a ground state (*p*_*A*_) and a sparsely populated state (*p*_*B*_ > *ca*. 0.5%) (53, 55).

We interrogated peptide binding kinetics and thermodynamics with ^15^N CPMG RD by adding a small amount of WT or P182L peptide to a solution of ^2^H,^15^N-labeled ACD and recording ^15^N dispersions, i.e. the effective ^15^N transverse relaxation rate (*R*_2,*eff*_) measured as a function of the number of 180°refocusing pulses (*ν* _*CPMG*_). With ca. 2% of the WT peptide-ACD complex present, large dispersions were observed for residues in the β4/β8 groove (Fig. 4A, B). Importantly, in the absence of peptide, the β4/β8 groove resonances did not yield significant dispersions (Fig. 4A, B), indicating that the observed effects are specific to the peptide-bound state. In addition, the magnitude of the dispersions depended on the amount of added peptide (Supplementary Fig. 4), which revealed that the conformational exchange event arises from an intermolecular binding reaction, *i*.*e*. the minor state involves a peptide-bound form of the ACD. Thus, the observed dispersions in the presence of ca. 2% of the ACD-peptide complex reflect chemical exchange between the peptide-free and peptide-bound states. Similar data were obtained for the P182L peptide-ACD complex (Fig. 4A, B), albeit with more peptide added to reach ca. 2% of the complex, as expected from the significantly elevated K_*d*_ (Fig. 3).

**Fig. 4.**
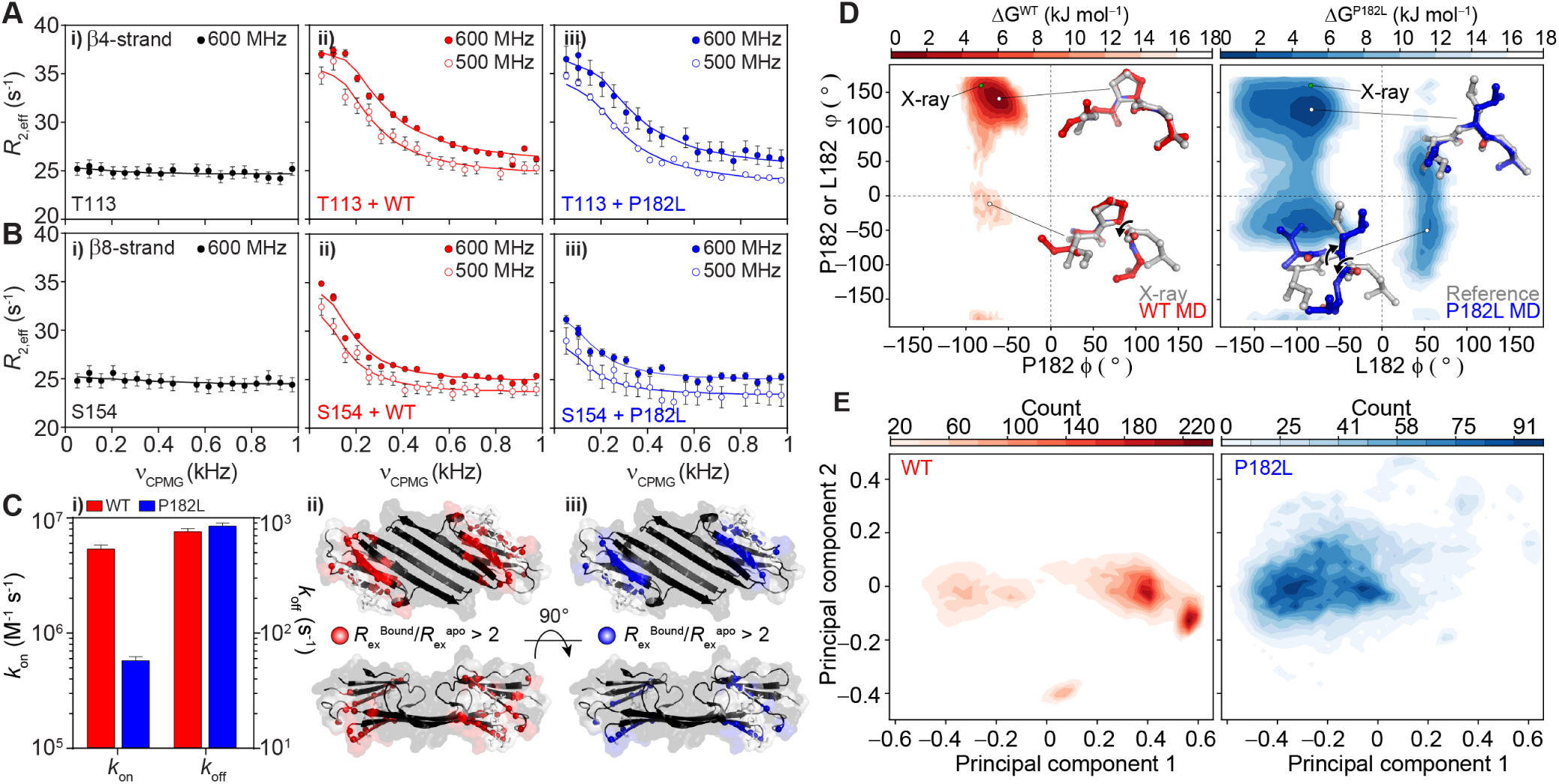
The P182L mutation perturbs IxI/V binding kinetics and conformational landscape.. (A, B) i. ^15^ N CPMG RD data for a residue in the β4 strand (T113) and the β8 strand (S154) in the absence of added peptide. CPMG data are shown for the same residues in the presence of ii. ca. 2% cHSP27-WT peptide complex or iii. ca. 2% of the cHSP27-P182L peptide complex. (**C**) i. The association (k_*on*_) and dissociation (k_*of f*_) rates for WT (red) and P182L (blue) IxI/V peptide binding to the ACD. Note that k_*of f*_ was calculated directly from the CPMG RD data (p_*A*_ k_*ex*_), whereas k_*on*_ was calculated using the K_*d*_ derived from the CSP titration and k_*off*_ from the CPMG RD analysis (see Methods, Supplementary Table 2) Residues that are impacted by the presence of a small amount of peptide are indicated by red (ii., WT) and blue (iii., P182L) spheres. In both cases, residues in similar ACD regions show enhanced R_*ex*_, indicative of chemical exchange at similar sites. (**D**) Free energy landscape of the WT (left) and P182L (right) peptides calculated from the MD trajectories as a function of the P182 or L182 *ϕ* and *ψ* angles. A lower value of ΔG corresponds to a higher relative population. Select IxI/V conformations extracted from the MD simulations are shown for the indicated *ϕ* and *ψ* angle pairs in red (WT MD) and blue (P182L MD). The conformation of the IxI/V motif in the crystal structure is shown in grey (X-ray) and the P182L reference structure in grey (Reference), as obtained with *in silico* mutagenesis using PyMol. The black arrows indicate rotations about the *ϕ* or *ψ* angles. The green circle denotes the *ϕ* and *ψ* angles of P182 when bound to cHSP27, measured from the crystal structure. (**E** Contour plot showing the first and second principal components of a PCA for the MD trajectories for the WT (left) and P182L peptide (right). The larger spread in the P182L peptide suggests a larger sampling of the conformational landscape.

CPMG RD data for the WT and P182L peptides were recorded at two static magnetic field strengths and analyzed using a two-state model of chemical exchange to extract kinetic and structural information. As observed earlier with CSPs, the CPMG RD data show that similar ACD regions experienced conformational exchange in both peptide-bound states (Fig. 4C ii., iii.), indicating that the WT and P182L peptides recognize and bind to the same site on the ACD (Supplementary Fig. 5, Supplementary Table 1). Moreover, the dissociation rates of the bound peptides from the ACD-peptide complexes were highly similar (Fig. 4C i.). We used the k_*off*_ values in conjunction with the K_*d*_ values we derived from CSPs to calculate the association rates (k_*on*_) for the two peptides, which differed by a factor of ca. 10 (Fig. 4C i.), indicating that the ca. 10-fold difference in binding affinity largely originates from an attenuated k_*on*_ rate for the P182L peptide binding. Note that, while both the CSP and CPMG RD data fit to a two-state model, using the CPMG RD data to directly calculate the K_*d*_ yields higher values than K_*d*_ values measured by CSP analysis for both WT and P182L peptides (Supplementary Table 2), likely reflective of the sensitivity of CPMG RD experiments to additional *µ*s-ms dynamics within the bound complex. However, the CPMG RD-derived K_*d*_ for the P182L peptide is 4-fold weaker than for WT peptide, with a 4-fold lower k_*on*_ value for the P182L peptide (Supplementary Table 2). Thus, both the CSP and CPMG RD data point to attenuated association of the P182L IxI/V motif, although they may be reporting on slightly different aspects of IxI/V binding.

### The P182L IxI/V motif samples a larger range of conformational space

We next sought to understand the mechanistic basis of these alterations. We hypothesized that, in its unbound form, the rigid Pro residue in the WT IxI/V motif would restrict overall flexibility of the adjacent Ile and Val residues and thereby ‘lock’ the IxI/V motif in place for subsequent binding to the ACD. By contrast, the P182L mutation would remove the steric hindrance of the restrictive Pro side-chain and enable easier exploration of the accessible conformational space.

To test these hypotheses, we performed 200 ns of all-atom molecular dynamics (MD) simulations on peptides of the same compositions used for the NMR binding experiments (Fig. 4D, E). The *ϕ* and *ψ* dihedral angles of P182 (WT) or L182 (P182L) directly impact on the conformation of the adjacent I181 and V183 residues within the IxI/V motif. We computed the free energy landscape for both the WT and P182L peptides as a function of *ϕ* and *ψ* dihedral angles for P182 or L182 (Fig. 4D). The analysis revealed that that central Pro residue in the WT IxI/V motif indeed significantly limits the sampling of backbone *ϕ* and *ψ* dihedral angles to a narrow region in Ramachandran space, located near –70°and +140°, respectively (Fig. 4D). Notably, the conformation of the bound state of the IxI/V motif. as determined by X-ray crystallography (20), exhibits similar P182 *ϕ*/*ψ* angles at –85°/+163°(Fig. 4D). In contrast, the P182L variant explores a much larger area of Ramachandran space, with highly populated conformers present near L182 *ϕ*/*ψ* dihedrals of –80°/+140°, –80°/–50°, +50°/–50°, and +50°/+50°(Fig. 4D).

The P182L conformations present in these free energy landscape valleys have significantly changed with respect to the conformation of the bound IxI/V motif (Fig. 4D). For example, the population of P182L conformers near *ϕ*/*ψ* +50°/-50°no longer adopt the necessary conformation for binding to the β4/β8 groove of the ACD: large rotations about the I181 CO–P182 N and P182 CO–V183 CA bonds have moved the side-chains of I181 and V183 out of plane (Fig. 4D), which would sterically clash with the atoms in the β4/β8 groove.

In addition to a comparison of the *ϕ*/*ψ* dihedral angles about the P182 or L182 residue, we performed a principal component analysis (PCA) over the MD trajectories to capture variations between the predominant structural and dynamical features of the two peptides. The conformations of the WT peptide predominantly occupy two clusters in the space of the first and second principal components (Fig. 4E), whereas the P182L variant instead shows a markedly broader distribution (Fig. 4E). The larger spread in the PC space of the P182L peptide reflects a broader sampling of conformational space (Fig. 4E). The IxI/V motif in the P182L variant is thus more flexible and less frequently adopts the conformation required to bind to the hydrophobic β4/β8 groove.

### Bioinformatics analyses identify HSP27-interacting proteins with exposed IxI/V motifs

Our results above demonstrate that the P182L mutation causes HSP27 to assemble into significantly larger oligomers by disrupting the binding of its own IxI/V motif. In addition, the P182L mutation also changes the affinities of its interactions with other proteins: the poly C-binding protein 1 (PCBP1) bound with higher affinity to the P182L variant of HSP27 (56). We note that PCBP1 contains numerous IxI/V motifs and therefore could compete with the IxI/V motif in HSP27 for binding to its β4/β8 groove (57). Since our data demonstrate that the IxI/V-binding site in P182L would be more accessible to IxI/V-containing proteins, we hypothesized that interactions between HSP27 and other IxI/V-containing proteins will be strengthened by the P182L mutation.

To test this hypothesis, we first searched the human proteome for all instances of the IxI/V SLiM according to its traditional definition of [V/I]X[V/I] and then determined how many of these proteins overlapped with previously identified HSP27-binding proteins. The human proteome search yielded a total of 128,604 instances or is 1.1% of all tripeptides (Supplementary Table 3, Supplementary Fig. 6). Next, we limited our search to the disordered regions of the proteome, since an IxI/V motif would need to be accessible to HSP27 and SLiMs typically occur in unstructured regions of proteins. This reduces the number of [V/I]X[V/I] motifs to 10,350 (Supplementary Table 3). Differences between the frequencies of [V/I]X[V/I] motifs in disordered and structured regions provides further insight into the amino acids that are enriched in disordered regions (Fig. 5A). The largest enrichment in disordered [V/I]X[V/I] motifs is found for X = proline (+7.1%), whereas the largest depletion is observed for X = leucine (–3.8%) (Fig. 5A). These results demonstrate that, in disordered regions, proline-containing [V/I]X[V/I] SLiMs are the most abundant type.

**Fig. 5.**
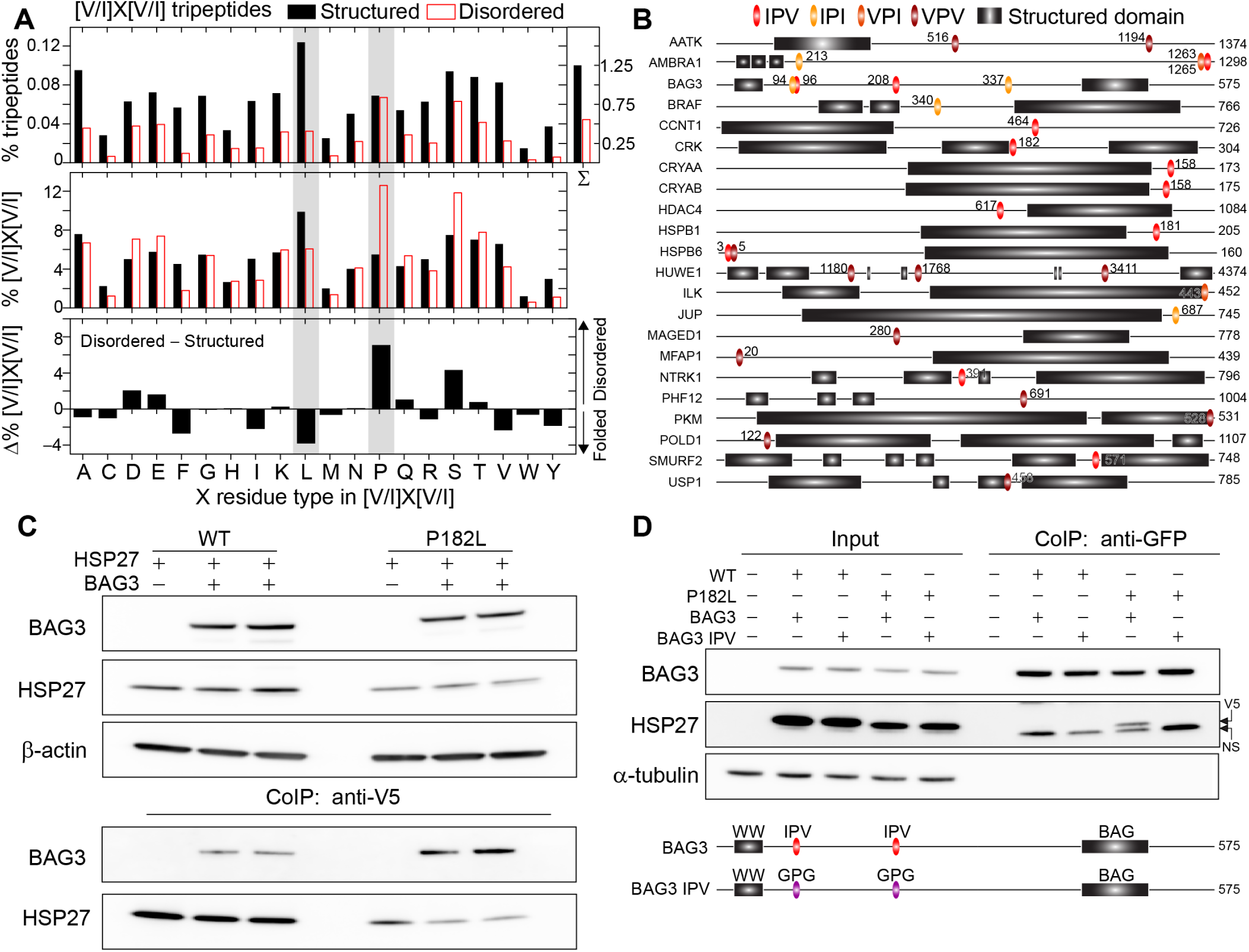
The P182L variant binds with enhanced affinity to an IxI/V-containing protein (BAG3) and bioinformatic analysis of IxI/V-containing proteins in the proteome. **(A)** Percentage of [V/I]X[V/I] tripeptides among (top) the total number of all tripeptides (% tripeptides) or (middle) the total number of [V/I]X[V/I] tripeptides in the structured (black) or disordered regions (red) ([V/I]X[V/I] %) shown as a function of the X residue type. (Bottom) The difference between [V/I]X[V/I] % for disordered and structured regions shows the enrichment (positive) or depletion (negative) of specific [V/I]X[V/I] motifs in disordered regions. The grey bars indicate the [V/I]L[V/I] and [V/I]P[V/I] motifs studied in this work, i.e. ILV and IPV. (**B**) Linear domain depiction of the 22 proteins that both interact with HSP27 and contain [V/I]X[V/I] motifs. The location of their [V/I]P[V/I] SLiMs and structured domains are indicated, along with their length. The numbers next to the [V/I]P[V/I] SLiMs indicate the starting residue number of the motif. (**C**) Co-immunoprecipitation using anti-V5 beads from HeLa cells stably overexpressing V5-epitope-tagged HSP27 (WT or P182L mutant) and transiently transfected with BAG3-eGFP. Western blots for anti-GFP (BAG3), V5 (HSPB1), and β-actin. (**D**) Co-immunoprecipitation using anti-GFP beads from HeLa cells stably overexpressing V5-epitope-tagged HSP27 (WT or P182L mutant) and transiently transfected with BAG3-eGFP (WT or IPV-mutant). Western blots for anti-GFP (BAG3), V5 (HSPB1), and α-tubulin. NS indicates a non-specific band..

We next sought to identify known HSP27-binding proteins that contain the [V/I]P[V/I] motif. We computed the intersection of known HSP27-binding proteins (58) and proteins that contain a [V/I]P[V/I] SLiM (Fig. 5B). Of the known 449 HSP27-binding proteins, a total of 22 proteins contain [V/I]P[V/I] SLiMs in disordered regions (Fig. 5B). Some of the identified proteins contain multiple such motifs in their primary sequences (Fig. 5B), which could further enhance their binding affinity to the P182L variant of HSP27 through multi-dentate interactions. Interestingly, five of the 22 proteins that we identified are implicated in stress granule formation (59, 60), and the 22 proteins in total make over 6,000 interactions with cellular proteins (58), suggesting that altered interactions with P182L could have wide-spread ramifications for other cellular processes. The HSP27-interacting [V/I]P[V/I]-containing proteins are listed in Supplementary Table 4 and span a diverse range of cellular functions, from chaperones and E3 ubiquitin ligases to kinases and regulatory proteins (Fig. 5B). A PANTHER overrepresentation test (61) indicates significant enrichment in multiple biological processes, including the negative regulation of cardiac and muscle cell apoptosis, regulation of neuronal apoptosis, the positive regulation of protein Ser/Thr kinase activity, and the positive regulation of angiogenesis (Supplementary Table 5).

### The P182L mutation enhances binding affinity for BAG3 through increased availability of the HSP27 IxI/V binding site

Having demonstrated that the IxI/V SLiM is abundant in the human proteome and that there are numerous IxI/V-containing proteins that interact with HSP27, we sought to investigate whether the P182L form of HSP27 binds with higher affinity to IxI/V-bearing proteins. To this end, we investigated in cells the binding of the P182L mutant to two established HSP27-interacting proteins with IxI/V motifs. The first was PCBP1, which carries multiple IxI/V motifs, and we previously showed to have increased affinity for the P182L mutant HSP27 protein (56). We substituted all six candidate IxI/V motifs by alanines (AxA); however, none of the substitutions yielded a stable protein in HeLa cells and therefore could not be used for further experiments. We therefore investigated a second candidate that contains an IxI/V motif, Bcl2-associated athanogene 3 (BAG3). This cochaperone contains two IxI/V motifs in its N-terminus (Fig. 5B, C), and previous work has indicated that BAG3 binds to sHSPs, including HSP27, via these IPV motifs (37, 62, 63). To assess whether BAG3 binds more tightly to the P182L mutant, we performed co-immunoprecipitation (coIP) experiments on HeLa cells stably overexpressing V5-epitope-tagged WT or P182L mutant HSP27 (Fig. 5C). Transient transfection of these cells with BAG3-eGFP demonstrated that BAG3 binds more tightly to the P182L mutant of HSP27 (Fig. 5C). Given the increased availability of the IxI/V binding site in the ACD of P182L HSP27, we hypothesized that mutating both IxI/V motifs in BAG3 would abrogate this interaction. Indeed, we created a BAG3 mutant in which the IPV motifs were mutated to GPG (BAG3-IPV mutant) and observed that it lost its interaction with HSP27 (Fig. 5D). These findings suggest that IxI/V-containing proteins bind more tightly to the P182L mutant due to the increased availability of the IxI/V binding site within this variant of HSP27.

## Discussion

We have investigated the biophysical and functional impact of the P182L mutation in the highly conserved IxI/V motif of HSP27. We found that the clinically severe P182L variant of HSP27 exists in a significantly larger oligomeric form, with an average mass *in vitro* that is nearly 30-fold higher than WT HSP27 (Fig. 2). In addition, sucrose gradient fractionation and expansion microscopy confirmed an increased oligomer mass inside cells (Fig. 2). Thus, the P182L mutant forms larger oligomeric structures in vitro and in vivo. Given the striking increase in the average mass of P182L oligomers, we assayed the ability of the P182L variant to protect against the aggregation of substrate proteins and found that it was devoid of chaperone activity (Fig. 1). This result is consistent with previous data on the P182S variant (40), although the P182L variant appears to form much larger oligomers. The relation between sHSP oligomer size and chaperone activity remains unclear, as it has been demonstrated that sHSP monomers are more active than dimers (44, 64) whereas larger oligomers of ABC can exhibit higher chaperone activity than the WT protein (65). To our knowledge, the P182L variant of HSP27 assembles into the largest soluble sHSP oligomers to-date, surpassing the mass of E. coli IbpB (2-3 MDa) (66). In addition, the P182L variant forms significantly larger oligomers than any of the CMT-related mutants of HSP27 studied thus far across the entire polypeptide sequence (38–40).

As the structural basis for the dysregulation of HSP27 by the P182L mutation has remained unknown, we sought to provide molecular details regarding the impact of this single amino acid mutation. The significance of this SLiM is indicated by its evolutionary conservation (Fig. 1), despite its occurrence in an IDR where frequently positional conservation is not observed (67, 68). We used CPMG RD (Fig. 3, Fig. 4) to quantify the kinetics of IxI/V binding to the ACD, and found that the P182L mutation drastically lowered the binding affinity for the ACD, largely through attenuation of the association rate (Fig. 4). Interestingly, the CPMG RD-derived dissociation rate (Fig. 4, Supplementary Table 1) and CSPs induced by peptide binding to the ACD (Fig. 3) were similar for both the WT and P182L forms. Furthermore, even though the resonances from residues in the β4/β8 groove could not be followed during the IxI/V titrations and therefore their CSPs could not be measured, CPMG RD enabled direct quantification of ^15^N |Δ*ω*| differences induced by IxI/V binding for these resonances, which were highly similar in both the WT and P182L peptide-bound forms (Supplementary Fig. 5). Together, the NMR results indicate that the P182L variant resembles the WT form when its IxI/V motifs are bound, but a lower association rate for the P182L peptide decreases the observed binding affinity. An explanation for this decreased k_*on*_ rate is provided by our MD simulations (Fig. 4), which show that Pro182 restricts the conformational landscape of the IxI/V motif, thereby placing it in a binding-competent conformation more often than the P182L variant. The larger sampling of conformational space by the P182L variant could thus contribute to its slower association rate.

If the P182L variant decreases the overall binding affinity for the IxI/V motif, then how does the protein form larger oligomers than the WT protein? Previous work on the similar sHSP ABC established that concomitant binding of two IxI/V motifs facilitates subunit exchange via ejection of a monomeric subunit from the oligomer (24, 25). Thus, were the IxI/V binding affinity to be lowered, subunit exchange would occur more slowly and the average oligomer size would increase (24, 25). Previous studies independently support such a mechanism, as other HSP27 mutants in the IxI/V motif have been shown to increase the molecular mass, including the HSP27 variants P182S (40) and GPG (34), and other IxI/V mutations (32–35). In addition, it has been demonstrated that other regions of sHSPs can also interact with the ACD, including NTD-β4/β8 groove interactions observed in a crystal structure of HSPB6 (69) and NTD-ACD dimer interface contacts as seen in a crystal structure of the HSPB2-HSPB3 hetero-tetramer (70). The CTRs of sHSPs play key roles in promoting solubility and contributing to the regulation of interactions with other proteins (71, 72), similar to the roles of other intrinsically disordered regions (73). In addition, the CTR of HSPB2 was shown to regulate its ability to undergo liquid-liquid phase separation (74). In addition, the CTR of HSPB2 was shown to regulate its ability to undergo liquid-liquid phase separation. Our data indicate that, alongside these other regulatory roles of the CTR, alteration of the binding properties of the IxI/V motif significantly affects the oligomeric landscape of HSP27.

In the full-length protein, the P182L IxI/V motif would be unbound more often than the WT form, and thus expose its dynamic CTR to solution more frequently than the WT protein. Less binding of the P182L IxI/V motif to the ACD would leave its β4/β8 groove more accessible to interactions with other parts of the protein (e.g. the NTD of itself or IxI/V of WT HSP27) or other proteins (37, 75, 76). Indeed, for a different IxI/V-containing protein (BAG3), our coIP data demonstrate that it binds more tightly to the P182L variant of HSP27 than the WT form, and that this interaction depends on the IPV motifs in BAG3 (Fig. 5). This co-chaperone/chaperone interaction promotes new protein-protein interactions, in part by bringing together HSP27 with other BAG3-bound chaperones, including HSP70 (37, 75). Moreover, BAG3 is recruited to stress granules via its interactions with HSPB8 where the BAG3-HSP8-HSP70 complex plays a key role in ensuring stress granule functionality (77). Disturbing the dynamic balance of the HSP27-BAG3 interaction might therefore affect a wider chaperone network and contribute to destabilization of proteostasis (78). Notably, mutations in the BAG3 IPV motifs also contribute to human disease: a Pro-to-Leu mutation is implicated in myofibrillar myopathy (34, 75, 79) and another Pro-to-Ser mutation is implicated in CMT disease (80). Thus, disruption of the IxI/V SLiMs in HSP27 and BAG3 contributes to the onset of pathogenic human diseases.

More generally, our coIP results suggest that HSP27-interacting proteins that carry IxI/V motifs may encounter reduced competition for the P182L IxI/V binding site and therefore bind to HSP27 more frequently. As a consequence, the interacting partner may become dysregulated due to aberrant interactions with HSP27 P182L that disrupt other interaction networks. Subtle changes to complex interaction networks can manifest in numerous human diseases (81), and so-called ‘hub’ proteins that facilitate a wide range of interactions are often mutated in diseases, such as cancers. A previous study found that competing interactions between HSP27 self-binding and substrate binding regulated HSP27 function and oligomerization (34). Our bioinformatics analysis identified 22 known HSP27 interactors that bear the IxI/V motif and are strong candidates to bind with higher affinity for the P182L variant. Based on data deposited in the BioGrid (58), these 22 proteins alone are known to make over 6,000 interactions with cellular proteins (Supplementary Table 4), in addition to numerous interactions with other biomolecules such as nucleic acids. Therefore, tighter interactions with P182L could disrupt processes that depend on transient interactions between these proteins or enzymes. In addition, our identification of [V/I]P[V/I] SLiM-containing proteins outlines a list of targets that could potentially bind to HSP27 via weak, transient interactions with its β4/β8 groove. In the crowded cellular environment, locally increased concentrations of available IxI/V motifs, either through compartmentalization or oligomerization, could facilitate interactions with HSP27.

The disrupted interaction network caused by the P182L mutation most strongly impacts WT HSP27 itself. Mutations in HSP27 that cause CMT disease and dHMN, including P182L, are primarily autosomal dominant, meaning that one allele will contain the mutated form of the gene and the other allele will contain the WT gene. Assuming relatively equal levels of expression in vivo, there will thus be a mixed population of WT and P182L HSP27 present inside the cell and hetero-oligomers of varying ratios of WT:P182L will presumably exist, as has been established for other disease-causing variants of HSP27 in vitro (82). Therefore, the available cellular pool of WT HSP27 will not only be depleted by having only one allele from which it is expressed, but also by the local sequestration of the WT protein by P182L oligomers. Interestingly, a recent NMR study of heterooligomerization between WT and mutated p97 implicated in an autosomal dominant neurological disease found evidence of allosteric communication between the protomers (83). It will thus be of interest to examine the role of inter-protomer interactions between WT and P182L forms of HSP27.

Similar to P182L, the S187L variant of HSP27 was found to aggregate in vivo (13), but it remains unclear if the aggregation is caused by increasing the hydrophobicity of the disordered CTR or if the mutation affects the binding affinity of the IxI/V motif. It has been proposed that the flexible CTR provides solubility to sHSP oligomers (71), so altering the hydrophobicity of this region may contribute to insolubility. Alternatively, lowering the affinity of the IxI/V motif for the ACD may allosterically trigger a change in the conformation of the NTD or its inter-subunit interactions. In addition, multiple lines of evidence point to the significance of the NTR in regulating oligomeric assembly (84), some of which contain IxI/V motifs or similar variations thereof (70), with oligomerization still observed for CTR-truncated forms of HSP27, but not NTR-truncated forms (20). In addition, it is interesting to note that mutations in other sHSPs, including α- and γ-crystallin, are linked to diseases such as cardiomyopathy and congenital cataract formation. Some of these disease-causing variants do not significantly change the three-dimensional structure of the protein (85–88), suggesting that subtle alterations to oligomeric assembly or other protein-protein interactions may play a role in disease onset.

In summary, our results provide an unexpected mechanistic explanation for the consequences of a mutation in the conserved IxI/V motif, which causes a severe form of CMT and dHMN. We observed that the P182L mutation significantly increases the average molecular mass of soluble HSP27 oligomers both in vitro and in vivo. We investigated the binding of the IxI/V motif to the ACD in detail with NMR spectroscopy and determined that the affinity is significantly attenuated in the P182L variant, with the K_*d*_ increased by nearly one order of magnitude. A consequence of the lowered binding affinity in the P182L variant manifests as a more available IxI/V binding site in the β4/β8 groove, which can also bind to IxI/V motifs present in other proteins. We observed that the IxI/V-containing protein BAG3 binds with increased affinity for the P182L variant in an IxI/V-dependent manner. In addition, bioinformatics analyses detected 22 known HSP27 interactors that harbor IxI/V motifs and are potential targets that could interact with increased affinity for the P182L variant, which could alter protein-protein interaction networks in vivo. Given the high level of conservation in the IxI/V motif across various mammalian sHSPs, we anticipate that our results contained herein will have learnings for other sHSP systems, and implicate this motif as a key mediator of sHSP-target interactions.

## Materials and Methods

### Proteins and peptides

Peptides were synthesized by Biomatik with N-terminal acetylation and C-terminal amidation. The peptides were dissolved in NMR buffer (30 mM sodium phosphate, 2 mM EDTA, pH 7), but the pH values of the solutions were found to be in the 4-5 range. As cHSP27 is highly pH-sensitive (64, 89), the pH of all peptide solutions was corrected to 7 using small volumes of concentrated NaOH.

Protein samples for NMR, chaperone activity assays, SEC-MALS, and negative-stain EM were expressed in E. coli and purified as described previously (64). WT HSP27 comprises residues 1–205, encompassing the entire amino acid sequence. The P182L mutation was introduced into the WT HSP27 expression plasmid via site-directed mutagenesis. The full-length P182L variant of HSP27 was expressed as the WT form but went into inclusion bodies. Inclusion bodies were solubilized in 8M urea, spun at 20,000 x g for 20 minutes, and then dialyzed into 30 mM sodium phosphate, 100 mM NaCl, 2 mM EDTA, pH 7 buffer. The solution was filtered with a 0.22 µm filter and then loaded onto a Superdex 200 gel filtration column equilibrated in 30 mM sodium phosphate buffer, 100 mM NaCl, 2 mM EDTA at pH 7.

cHSP27 encompasses residues 84–171 of HSP27, and the expression plasmid contains an N-terminal hexahistidine tag followed by a tobacco etch virus (TEV) protease cleavage site. The Gly overhang following TEV protease cleavage corresponds to G84 in the HSP27 amino acid sequence. cHSP27 was expressed and purified as described previously (64). Following exchange into a buffer without reducing agent, cHSP27 forms an intermolecular disulfide bond involving C137 from adjacent subunits. For NMR titration and relaxation dispersion studies, this disulfide bond was left intact to minimize contributions from exchange between the monomer and dimer. The formation of the disulfide bond is readily identifiable in the 2D^1^*H*-15 *N* HSQC spectrum of 15*N* -cHSP27 via the resonances from R136 and C137; upon reduction, these resonances become very weak or broaden into the noise (64, 65). In addition, the oxidized dimer simplifies the analysis of titration and CPMG RD data, as only a single state of the protein is present, rather than a mixture of two forms (non-covalent dimer and monomer).

Additional information concerning the biochemical and biophysical analyses, cell lines and cell culture, image processing and analysis, and bioinformatics analyses is provided in the SI Appendix.

## ACKNOWLEDGEMENTS

We are grateful to Heath Ecroyd (University of Wollongong) for the HSP27 expression plasmid, and we thank Katrien Spaas of the VIB CMN Neuromics Support Facility and Vicky De Winter for assistance in expansion microscopy sample preparation and cloning, respectively. This research was supported in part by the Research Foundation - Flanders (FWO-Vlaanderen; to V.T.), the Medical Foundation Queen Elisabeth (GSKE; to V.T.), the Association Belge contre les Maladies Neuromusculaires (ABMM; to E.A. and V.T.), the Rotary ‘Hope in Head’ program (to E.A. and V.T.) and the H2020 Solve-RD program ‘Solving the unsolved rare diseases’ under grant agreement 2017-779 257 (to V.T.). T.R.A acknowledges funding from the NIDDK and the NIH Oxford-Cambridge Scholars Program. A.J.B. holds a David Phillips Fellowship from the Biotechnology and Biosciences Research Council (BB/J014346/1). J.L.P.B. thanks the Engineering and Physical Sciences Research Council (EP/J01835X/1) and Biotechnology and Biosciences Research Council (BB/J018082/1).

## Supplementary Information

### Chaperone activity assays

Chaperone activity assays were completed in triplicate, with the mean and one standard deviation reported. Assays were performed using a 96-well plate and a FLUOstar Omega Microplate Reader or Tecan Infinite M200 PRO plate reader. The substrates were porcine heart MDH (Sigma-Aldrich), which was prepared at 0.2 µM and was incubated at 40 °C in 30 mM sodium phosphate, 100 mM NaCl, 2 mM EDTA at pH 7. The increase in absorbance at 340 nm was monitored over time in the absence and presence of 0.5 µM WT or P182L HSP27. The MDH aggregation assay proceeded for 3 hours. Likewise, the aggregation of 40 µM human insulin (Sigma-Aldrich) at 40 °C in the same buffer as above was monitored by the increase in absorbance at 340 nm upon the addition of 1 mM DTT. Insulin aggregation reactions were monitored in the absence or presence of 20 µM WT or P182L HSP27, and the reactions proceeded for 0. 5 hours. Control experiments were performed using 20 µM WT or P182L HSP27 incubated alone in the same buffer as above at 40 °C for 3 hours. Chaperone activity (Supplementary Fig. 1) is defined as 1 – Θ, where Θ is the maximum intensity for the chaperone plus substrate mixture divided by the maximum intensity of the substrate alone. Values of 1 and 0 respectively indicate maximum protection and the absence of protein against aggregation.

### SEC-MALS

SMolecular masses were estimated by analytical SEC with in-line MALS (DAWN Heleos-II, Wyatt Technology Inc., Santa Barbara, CA), refractive index (Optilab T-rEX, Wyatt Technology, Inc.), and UV detectors (Waters 2487, Waters Corp., Milford, MA), along with ASTRA version 6 software that was provided with the instrument. In all cases, 250 µg of protein was injected to a pre-equilibrated Superdex-200 column (for P182L and HSP27) or Superdex-75 column (for cHSP27 with the C137S mutation) in 30 mM sodium phosphate, 100 mM NaCl, 2 mM EDTA buffer at pH 7 and eluted at a flow rate of 0.5 mL/min. The monomer concentration of WT and P182L HSP27 corresponded to 40 µM. The calculated molar masses of HSP27, P182L, and cHSP27(C137S) were 470 kDa, 13600 kDa, and 17.5 kDa, respectively, within 1-1.2% error. The theoretical mass of the cHSP27(C137S) dimer is 19.8 kDa, so the slightly lower value as determined by SEC-MALS likely reflects contributions from exchange with the free monomer. Note that the C137S mutation does not impact the overall structure of the cHSP27 dimer (64). The molecular mass of WT HSP27 is in good agreement with previous publications (84, 90), with the former reference indicating 670 kDa by analytical SEC and the latter 400 kDa by SEC-MALS.

### Negative-stain EM

Samples for negative-stain EM were prepared at 4.5 µM monomer concentration in 30 mM sodium phosphate, 100 mM NaCl, 2 mM EDTA at pH 7. Samples were loaded onto carbon-coated copper EM grids (Ultrathin Carbon Film/Holey Carbon; Ted Pella) and incubated there for 60 seconds. Excess sample was blotted off the grid, and then the grids were washed with deionized water and stained with 0.5% uranyl formate for 20 seconds. Excess stain was removed by blotting. Negative-stain EM images were then collected on a FEI Tecnai T12 electron microscope operating at 120 kV, which was equipped with a Gatan US1000 CCD camera.

### Cell lines, cell culture, and transfection

HeLa cells were cultured in MEM medium (ThermoFisher Scientific) supplemented with 10% fetal bovine serum (FBS; ThermoFisher Scientific), 1% glutamine (ThermoFisher Scientific), and 1% penicillin/streptomycin. Cells were maintained at 37 °C and 5% CO2 atmosphere.

To create HSP27 CRISPR knockout lines, we generated a sgRNA containing pSpCas9(BB)-2A-PURO (PX459) V2.0 vector (Addgene, 62988). The most specific sgRNA was selected with crispr.mit.edu by targeting the first exon of the HSP27 gene. Through molecular cloning following Ran et al. (91), the sgRNA was cloned into the BbsI restriction site and successful insertion was verified by Sanger sequencing. HeLa cells were seeded in a 6-well plate and transiently transfected using polyethylenimine (PEI). After 24 hours, the medium was supplemented by 1 µg/ml puromycin to select for positively transfected cells. The puromycin was removed after 72 hours as all the cells were dead in a non-transfected well. The positively transfected cells were then serially diluted in a 96-well plate with an estimated 0.5 cells/well. Wells that contained clonal expansion for more than one colony were discarded. Other wells starting from a single colony, were expanded and collected to verify the genome editing both at the protein and DNA level. Knockout clones were identified by western blotting using a monoclonal HSP27 antibody (Enzo Life Sciences SPA-800). The successful clones were screened further by DNA sequencing of the edited genomic region, after subcloning the amplicons into pUC19 plasmids. This allowed to identify the genome edits for each of the respective alleles. Only clones with an identified stop codon on each allele were selected for further experiments.

For microscopy experiments, HSP27 knockout HeLa cells were grown on standard cover glasses (12 mm 1.5) at a concentration of 50.000 cells/well. The cells were transiently transfected with HSP27 wild-type or P182L mutant ORF in pLenti6/V5 plasmids (ThermoFisher Scientific).

### Expansion microscopy sample preparation and imaging

Cells were immunostained and processed for expansion microscopy employing a protocol by Chozinski et al. (92).Twenty-four hours after transfection, cells were fixed for 20 minutes in 3.2% paraformaldehyde and 0.1% glutaraldehyde in PBS with 5 minute reduction in borohydride. After three PBS washes, samples were blocked and permeabilized for 1 hour at room temperature in normal goat serum (1:500, Dako) and 0.5% Triton-X, dissolved in PBT buffer (PBS + 0.5% BSA + 0.02% Triton-X). Primary antibody incubation over night at 4 °C with 1:500 anti-HSP27 (Enzo Life Sciences SPA-800) in PBT was followed by four 5-minute and one 30-minute PBT washes, 1 hour secondary antibody incubation at room temperature (1:500 in PBT, AlexaFluor488 goat anti-mouse, ThermoFisher Scientific A11001), four 5-minute PBT washes and one 30-minute PBS wash. Control coverslips for conventional confocal microscopy were nuclear counter stained with Hoechst 33342 (1/20000 for 10 minutes at room temperature) and mounted on microscopy slides in antifade mounting medium (Dako). Samples for expansion microscopy were crosslinked for 10 minutes in 0.25% glutaraldehyde in PBS. Gelation was done in a mixture of 2 M NaCl, 2.5% (w/w) acrylamide, 0.15% (w/w) N,N’-methylenebisacrylamide, 8.625% (w/w) sodium acrylate in PBS with polymerization initiated with TEMED and APS. Polymerized gels were incubated for 30 minutes at 37 °C in a digestion buffer containing 8 U/ml proteinase K. Cover glasses were removed from the digested gels, which were placed in high volumes (>30 mL) of distilled water that were exchanged at least 5 times until full expansion of the gels. Finally, the gels that had expanded 4.3 times in all dimensions were trimmed, nuclear counter stained with Hoechst 33342 (1/5000 in water for 30 minutes at room temperature), positioned in 50mm diameter glass bottom dishes (WillCo Wells GWSt-5040) and immobilized using 2% agarose.

Image stacks of the expanded sample were acquired on a Zeiss LSM700 with Plan-Apochromat 63×/1.40 objective, using 85nm × 85nm × 445 nm voxel dimensions (corresponding to approximately 20nm × 20nm × 100nm in a non-expanded sample). We measured minimal spot diameters (FWHM in expansion corrected scales) in the range of 60-80 nm (data not shown), confirming that the physical size of the particles is still below this limit and cannot be directly quantified.

### Image processing and analysis

Cytoplasmic regions of interest were randomly selected from image stacks (200 x 200 pixels, single slice) and the HSP27 distribution was analyzed using the Fiji distribution of ImageJ [83,84]. After noise filtering (Gaussian blur, sigma 0.7), high intensity spots were detected with the Find Maxima tool (fixed noise level 50). The intensity and size of the spots was measured with the GaussFitOnSpot plugin (Peter Haub and Tobias Meckel) using circle shape and Levenberg Marquard fit mode and 10 pixel (850 nm) rectangle half size. All images were processed in batch using ImageJ macro scripts. Data analysis and plotting was done using R [85]. Misfitted spots were removed by excluding spots smaller than 20 nm and larger than 200 nm (FWHM in expansion microscopy corrected dimensions). For each genotype, 15 cytoplasmic regions of the same size as the expansion microscopy images in Fig. 3C were analyzed (4 µm x 4 µm), which resulted in the detection of 3342 spots in total.

### Sucrose gradient assay

HeLa cells were lentivirally transduced with HSPB1-V5 (wild type or P182L mutant) and collected for protein extraction with RIPA buffer [1% Nonidet P-40, 150 mM NaCl, 0.1% SDS, 0.5% deoxycholic acid, 1 mM EDTA, 50 mM Tris-HCl pH 7.5, cOmplete Protease Inhibitor Cocktail (Roche Applied Science, Indianapolis, IN, USA), Phospho-STOP inhibitor mix (05 892 970 001, Roche Applied Science, Indianapolis, IN, USA)] for 30 min on ice. Protein concentrations were quantified using BCA (23225, Pierce BCA Protein Assay Kit). The sucrose gradient was generated by diluting a 80% sucrose stock solution in PBS to 70%, 60%, 50%, 40%, 30%, 20% and 10%. Equal amounts from each fraction were layered on top of each other and allowed to equilibrate on ice for 30 min. Equal amounts of protein lysate were then loaded on top of the gradient. Note that this is total whole cell protein lysate and thus contains both soluble and non-soluble proteins. Samples were centrifuged for 60 min at 55 000 rpm in a TLA-110 rotor (Beckman). Nine samples were collected and prepared for SDS-PAGE analysis. After separation on 4-12% NuPAGE gels (Life Technologies, Carlsbad, CA, USA). Proteins were transferred to nitrocellulose membranes (Hybond-P; GE Healthcare, Wauwatosa, WI, USA) and decorated with antibodies against V5 (R96025, Invitrogen, Carlsbad, CA, USA). Samples were detected using enhanced chemiluminescent ECL Plus (Pierce, Life Technologies, Carlsbad, CA, USA) and LAS4000 (GE Healthcare, Wauwatosa, WI, USA).

### Co-immunoprecipitation

Stable HeLa cell lines for HSPB1-V5 (wild type or P182L mutant) were transiently transfected with different wild type or IPV-mutant BAG3-GFP constructs using linear polyethylenimime (PEI) (PolySciences Europe, Hirschberg an der Bergstrasse, Germany). The IPV-mutant of BAG3-GFP was generated by site-directed mutagenesis by replacing both the first and second IPV-motif of BAG3 with GPG (the respective amino acids are 96-98 and 208-210). Twenty four hours after transfection, cells were lysed in lysis buffer [20mM Tris-HCl pH 7.4, 2.5 mM MgCl_2_, 100 mM KCl, 0.5% Nonidet P-40, Complete Protease inhibitor (Roche Applied Science, Indianapolis, IN, USA)] and incubated on ice for 30 min. Protein lysates were cleared by centrifugation for 10 min at 20,000 g and equal amounts of supernatant (NP40-soluble fraction only) was loaded on GFP-Trap beads (gta-100, Chromotek, Martinsried, Germany) or anti-V5 agarose beads (A7345, Sigma where [L]_*t*_ and [P]_*t*_ refer to the total concentration of ligand (peptide) and protein (cHSP27), respectively. Thus, at 80 µM of added WT peptide in the presence of 1.5 mM protein for a K_*d*_ of 125 µM, approximately 4.9% of cHSP27 should be bound. For the P182L peptide at a concentration of 200 µM in the presence of 1.5 mM protein for a K_*d*_ of 1200 µM, approximately 7.2% of cHSP27 should be bound. These calculations yield significantly larger values of *p*_*B*_ than what was experimentally measured by CPMG RD: 2.0% and 1.7% for WT and P182L, respectively. See below for a possible discussion on why.

Dispersions were fit to a model of two-state chemical exchange using the CATIA software (95). A total of 10 residues each from the WT and P182L peptide datasets were included in the global fit: T110, K112, T113, K114, T151, V153, S154, S156, L157, and S158 (Supplementary Table 1). These residues were selected based on a large increase in Rex in the presence of peptide as compared to in the absence of peptide. In oxidized cHSP27, residues D129, E130, R136, C137, and F138 also show dispersions, but these residues were not included in the fit, as they arise from non-peptide-binding related processes (64). Between the two CPMG RD datasets derived from WT and P182L peptides, a large degree of similarity in the fitted 15N |,Δω| values is observed (Supplementary Fig. 3), with an RMSD of 0.27 ppm. Since the |, Δω| values reflect the chemical shift changes upon binding to peptide, this agreement between the two independently fit datasets further highlights the similarity of the conformations of the peptide-bound states.

### Two- or three-state binding model

While both the CSP and CPMG RD data fit well to a model of two-state binding, the potential presence of a third-state is evident from both the discrepancy between *p*_*B*_ calculated from theory (eqn 5) and that measured by CPMG RD. For the CPMG RD data, the measured *p*_*B*_ values are lower than the expected values given the *K*_*d*_ values and the concentrations of ligand and protein. Using the CPMG RD-derived values of *k*_*off*_ and *k*_*on*_ yields *K*_*d*_ values that are significantly higher than those measured by CSPs: namely, for the WT peptide the CPMG-derived *K*_*d*_ is 2.51 mM and for the P182L peptide the *K*_*d*_ is 10.2 mM. The respective *k*_*off*_ and *k*_*on*_ values for WT (P182L) are 759 s^−1^ (840 s^−1^) and 3.02 × 10^5^ M^−1^ s^−1^ (8.26 × 10^4^ M^−1^ s^−1^). To convert the fitted *k*_*ex*_ values into *k*_*on*_ and *k*_*off*_, and calculate the *K*_*d*_, the following equations were used:

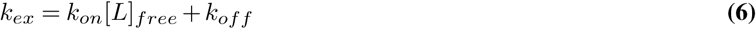

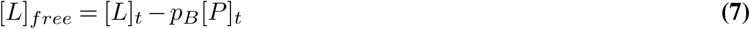

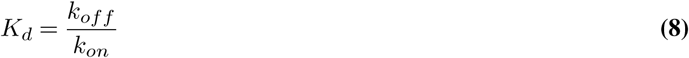

This discrepancy in binding affinity as measured by CSPs and CPMG RD could arise from a third state. The source of the broadening and lower than expected *p*_*B*_ values could be significant heterogeneity of the peptide when bound to cHSP27, leading to multiple interconverting states on the millisecond timescale. However, the fact that both CSP and CPMG RD data are well-fit by a model of two-state chemical exchange thus implies that the third state rapidly interconverts with one of the two states probed here. A potential source of broadening could arise from transient oligomerization. We recorded ^15^*N* relaxation in both the absence and presence of near-saturating peptide (95% bound), and the NMR-visible state has not changed its oligomeric state (Supplementary Fig. 7). The fitted *τ*_*c*_ values from ^15^*N T*_1_/*T*_2_ ratios (96) in both cases are consistent with a dimer of 20 kDa in mass at 310 K (*ca*. 10 ns). As such, we speculate that multiple conformations of the peptide-bound form may exist, with significant conformational interconversion on the microsecond timescale. Therefore, a plausible binding model may look like one of the scheme below:

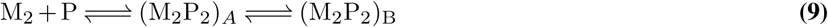

where the two different bound forms indicated inside parentheses rapidly interconvert on a microsecond timescale, which is similar to a previous NMR report (97).

### Molecular dynamics simulations

All-atom MD simulations of peptides containing the IxI/V motif and the P182L variant were performed using the Gromacs version 4.5.3 simulation package (98) with the AMBER99SB-ILDN force field and the TIP3P water model. To obtain a starting conformation of the IxI/V-containing peptide, chain B comprising residues I179-E186 of HSP27 was extracted from the crystal structure of the ACD bound to its IxI/V motif (PDB: 4mjh, (22)). The missing atoms were added with the Modeller web server (99). The P182L variant of the peptide was obtained by *in silico* mutagenesis of the P182 residue using PyMOL. After adding the missing atoms, including hydrogens, the WT and P182L peptides respectively contained 134 and 139 atoms.

Each MD simulation was performed in a rhombic dodecahedron of size 5.30 by 5.30 by 3.75 nm, which contained 3403 (WT) or 3401 (P182L) water molecules and one sodium ion to maintain charge neutrality. A total simulation time of 200 ns was achieved using a time step of 2 fs with 107 total steps. The temperature was maintained at 37 °C using the velocity-rescaling thermostat (100) and the average isotropic pressure was kept at 1 bar with the Parrinello-Rahman barostat (101). Long-range electrostatics were calculated with the Particle Mesh Ewald summation. Both peptides were subsequently energy minimized and equilibrated for 20 ns prior to analysis. Quantitative analyses of the trajectories including dihedral angle calculations and PCAs were performed using in-house Python scripts and the MDTraj Python package (102). The PCA was performed by combining the WT and P182L trajectories into a single trajectory, extracting only the Cα atoms, and aligning all frames to the crystal structure of the bound form of the WT IxI/V motif (PDB: 4mjh). The first two principal components were then calculated from this combined trajectory, with the residues at the N- and C-termini excluded from the analysis.

### Bioinformatic analyses

[V/I]P[V/I] motifs were quantified in the reference human proteome that contained only the longest transcript of each gene, with the UniProt Proteome ID UP000005640. Within the proteome, we identified predicted regions of intrinsic disorder comprising 20 residues or longer by running the DISOPRED3 software (103) on each gene and selecting contiguous regions with a disordered score of 0.7 or higher. An in-house Python script making use of the Biopython package (104) was written to search for and count specific peptide motifs within the proteome (M_*proteome*_) and the database of disordered regions (M_*dis*_). To obtain the total count of tripeptides, a sliding window of three amino acids was incremented over the proteome and disordered regions. To count specific tripeptide motifs [V/I]X[V/I] in structured regions of the proteome (M_*struct*_), we subtracted M_*dis*_ from M_*proteome*_ for each queried tripeptide j (1, …, N) in which the central position was iterated over each amino acid i (1, …, 20), as such:

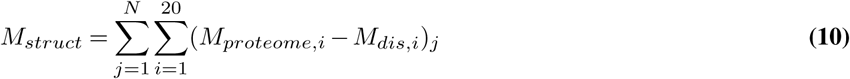

Removing the summation over *i* in equation 10 yields the count of specified tripeptides that include a fixed amino acid at the central position, e.g. [V/I]P[V/I]. To calculate the expected number of peptide motifs found in each database, we used the following equation:

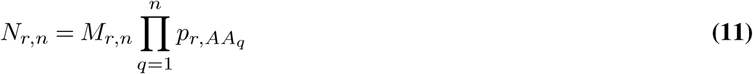

Where *p*_*r,AA*_ is the frequency of amino acid *AA* appearing in the database *r* (*r* = [proteome, structured, disordered]) and *M*_*r,n*_ is the total number of peptides of length *n* in the database *r*. The product sum is calculated by computing *p*_*r,AA*_ iteratively for each position q in the queried motif (*q* = 1, 2, …, *n*). This yields *N*_*r,n*_, the expected number of peptides of length n in the database *r*. This assumes that each amino acid in the queried peptide motif has an independent probability of appearing next to any other amino acid, and therefore we do not account for selective pressures imparted by, e.g. co-evolution of specific motifs. As an illustrative example, the queried tripeptide ISV within the database of disordered regions (*M*_*disordered*,3_ = 1,941,166) with frequencies of Ile, Ser, and Val corresponding to *p*_*disordered,Ile*_ = 0.0243, *p*_*disordered,Ser*_ = 0.1223, *p*_*disordered,V al*_ = 0.0446, yields an expected total of 257 ISV motifs. This value agrees closely with the observed number of ISV motifs, 299. The Python code that was used for these analyses and the list of disordered regions longer than 20 residues are available upon request.

For the 22 proteins found to interact with HSP27 and contain the [V/I]P[V/I] SLiM, we performed a PANTHER Overrepresentation Test (released 20190701) against the GO Ontology database (released 20190202), using the Homo sapiens reference list and the “GO biological process complete” annotation data set. Bonferroni correction for multiple testing was employed with a P value cut-off of 0.05 to filter the results.

---

**Supplementary Table 1.**
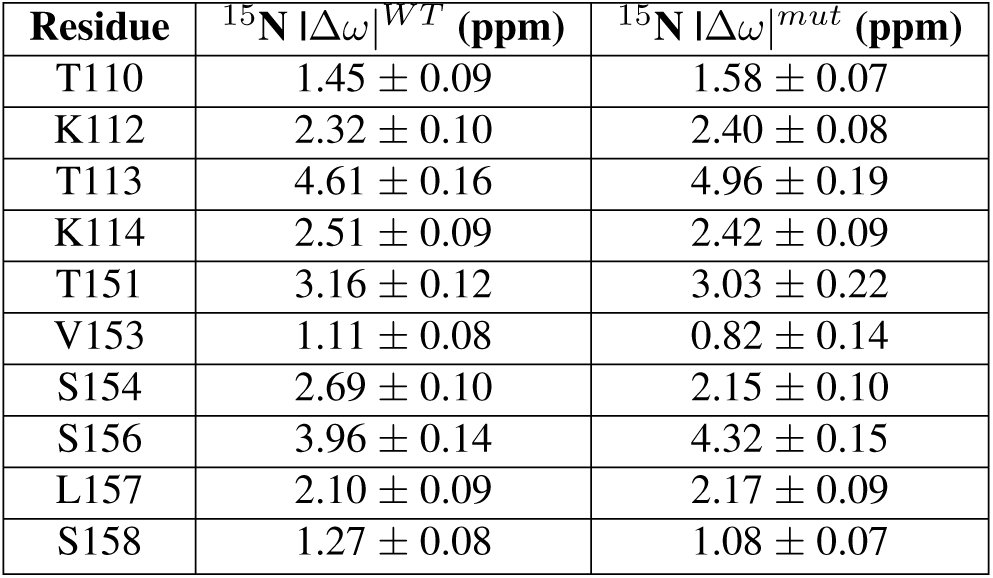
CPMG RD-derived ^15^N |Δω| values for WT- and P182L-bound cHSP27. ^15^N |Δω| values in cHSP27 derived from CPMG RD analysis in the presence of a small amount of WT or P182L peptide. Dispersions at 298 K at 500 and 600 MHz were included in the global fit; WT and P182L datasets were analyzed independently. The superscripts WT and mut refer to values obtained in the presence of WT peptide and P182L peptide, respectively. Both the fitted value and error are listed for each residue.

| **Residue** | **$^{15}\text{N}$ $\vert \Delta\omega\vert ^{WT}$ (ppm)** | **$^{15}\text{N}$ $\vert \Delta\omega\vert ^{mut}$ (ppm)** |
| --- | --- | --- |
| T110 | $1.45 \pm 0.09$ | $1.58 \pm 0.07$ |
| K112 | $2.32 \pm 0.10$ | $2.40 \pm 0.08$ |
| T113 | $4.61 \pm 0.16$ | $4.96 \pm 0.19$ |
| K114 | $2.51 \pm 0.09$ | $2.42 \pm 0.09$ |
| T151 | $3.16 \pm 0.12$ | $3.03 \pm 0.22$ |
| V153 | $1.11 \pm 0.08$ | $0.82 \pm 0.14$ |
| S154 | $2.69 \pm 0.10$ | $2.15 \pm 0.10$ |
| S156 | $3.96 \pm 0.14$ | $4.32 \pm 0.15$ |
| L157 | $2.10 \pm 0.09$ | $2.17 \pm 0.09$ |
| S158 | $1.27 \pm 0.08$ | $1.08 \pm 0.07$ |

**Supplementary Table 2.**
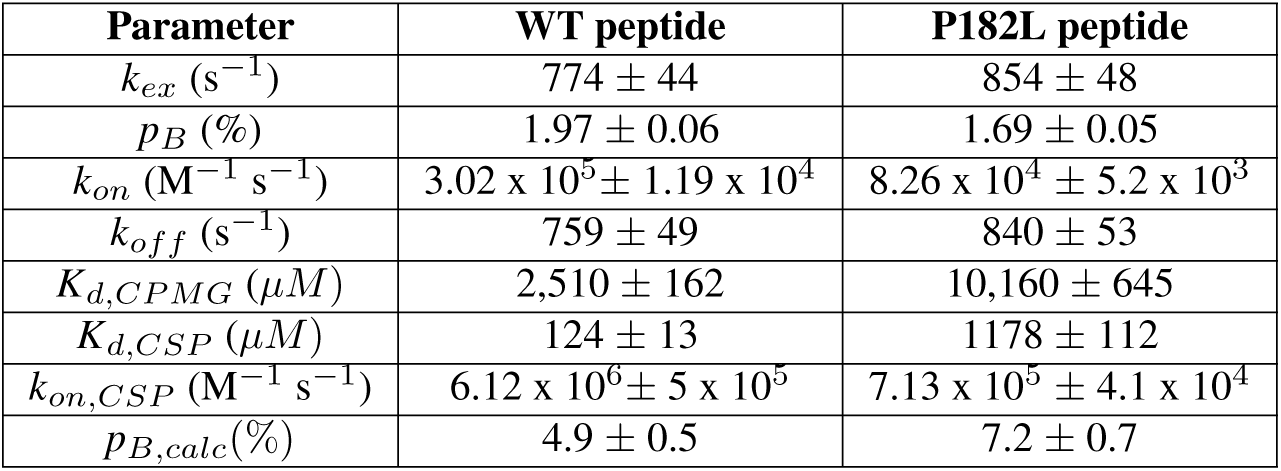
Kinetic parameters for WT and P182L peptide binding to cHSP27. Values of k_*on*_ (M^−1^ s^−1^), koff (s-1), and Kd (µM) as derived from CPMG RD studies on sparsely peptide-bound cHSP27. The Kd refers to the peptide-cHSP27 equilibrium, with koff and kon the dissociation and association rates, respectively. The superscripts CPMG and CSP respectively refer to values determined using CPMG RD or NMR titrations (CSP). The superscript calc,CSP refers to the calculated value based on the Kd determined from the CSP analysis.

**Supplementary Table 3.** Bioinformatics analyses of the IxI/V motif. The percentages shown in the final column were calculated with respect to ^*a*^ the total number of tripeptides in the listed databases or ^*b*^ the total number of [V/I]X[V/I] tripeptides. The number of tripeptides was calculated using a sliding window over each gene in the respective databases. The database “Disordered” includes both intrinsically disordered proteins and intrinsically disordered regions longer than 20 residues, as determined by a DISOPRED3 score larger than 0.7 (see Methods).

| Motif | Database | Count | % |
| --- | --- | --- | --- |
| XXX | Proteome | 11,373,813 | - |
|  | Structured | 9,432,647 | - |
|  | Disordered | 1,941,166 | - |
| [V/I]X[V/I] <sup>a</sup> | Proteome | 128,604 | 1.11% |
|  | Structured | 118,254 | 1.25% |
|  | Disordered | 10,350 | 0.53% |
| [V/I]P[V/I] <sup>b</sup> | Proteome | 7,803 | 6.07% |
|  | Structured | 6,500 | 5.50% |
|  | Disordered | 1,303 | 12.60% |
| [V/I]L[V/I] <sup>b</sup> | Proteome | 12,309 | 9.57% |
|  | Structured | 11,681 | 9.88% |
|  | Disordered | 628 | 6.07% |

**Supplementary Table 4.**
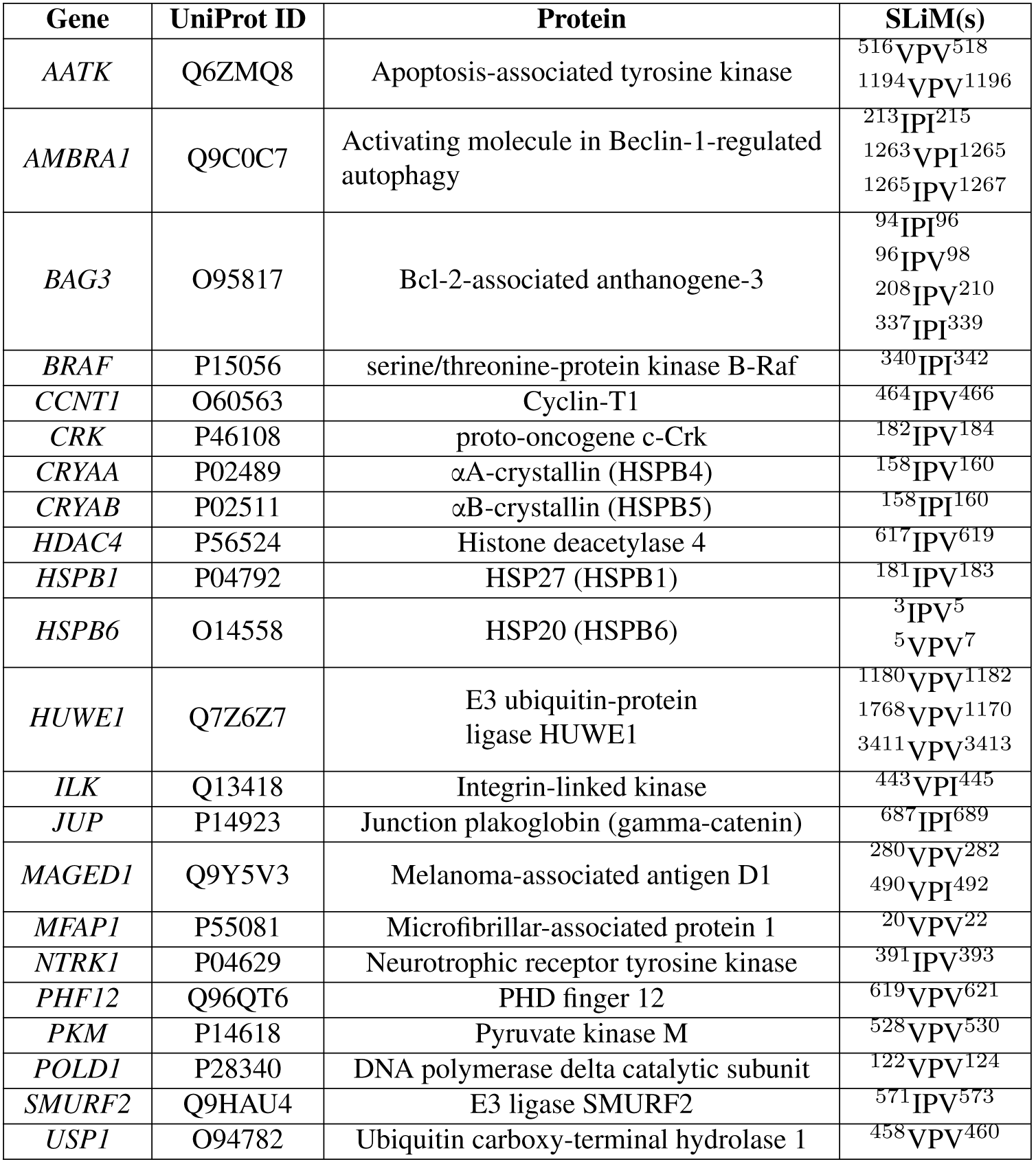
Known HSP27-interacting proteins that also contain IxI/V motifs. For each hit, the gene name, UniProt ID, protein name, and [V/I]P[V/I] SLiM composition and numbering are listed.

**Supplementary Table 5.** Pathways involving HSP27-interacting proteins with IxI/V motifs. The proteins listed in Supplementary Table 4 were subjected to a PANTHER overrepresentation test using the human proteome as a reference. Statistically significant biological processes are listed below (GO Biological Processes) with the total number of proteins in the reference proteome for the listed process (Proteome), the observed number of proteins in the queried dataset (Observed), the expected number of proteins given the size of the queried dataset (Expected), the fold enrichment over the expected value, and the P value. Bonferroni correction was used in the determination of statistical significance.

| GO Process | Proteome | Observed | Expected | Fold enrichment | P value |
| --- | --- | --- | --- | --- | --- |
| Negative regulation of cardiac muscle cell apoptotic process | 21 | 3 | 0.02 | 100 | 0.0175 |
| Negative regulation of intracellular transport | 64 | 4 | 0.07 | 59.65 | 0.00617 |
| Positive regulation of angiogenesis | 165 | 5 | 0.17 | 28.92 | 0.00676 |
| Regulation of neuron apoptotic process | 209 | 5 | 0.22 | 22.83 | 0.0202 |
| Positive regulation of protein Ser/Thr kinase activity | 334 | 6 | 0.35 | 17.14 | 0.00904 |
| Response to abiotic stimulus | 1153 | 9 | 1.21 | 7.45 | 0.0106 |

**Supplementary Figure 1.**
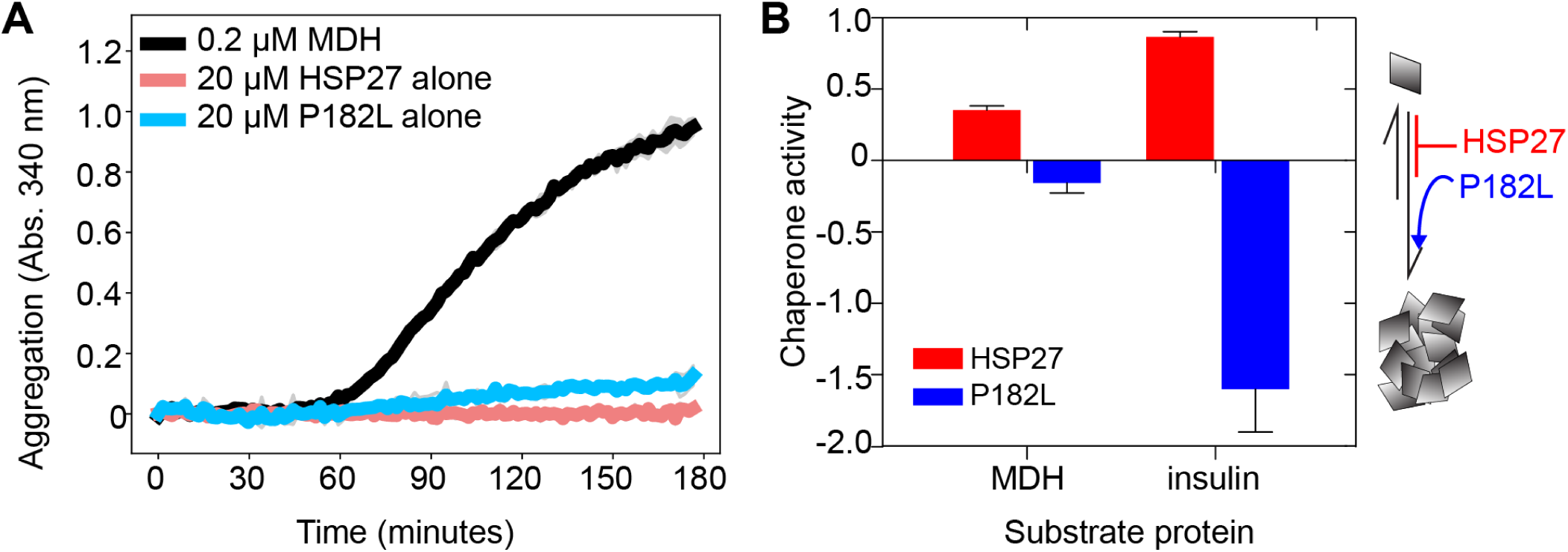
Control experiments for P182L and HSP27 self-aggregation. (**A**) Data from Figure 2 depicting the aggregation of 0.2 µM malate dehydrogenase (MDH, black) at 40 °C in the presence and absence of 0.5 µM HSP27 (red) or 0.5 µM P182L (blue). Alongside these experiments, controls were performed in which 20 µM of HSP27 (light red) and 20 µM of P182L (light blue) were incubated at 40 °C. There is no absorbance at 340 nm (Abs. 340 nm) for 20 µM HSP27, indicating the absence of aggregation, and only minor signals for 20 µM P182L. The buffer conditions were 30 mM sodium phosphate, 100 mM NaCl, 2 mM EDTA at pH 7. (**B**) The chaperone activity of HSP27 (red) and the P182L variant (blue) against MDH and insulin. Chaperone activity is defined as 1 –, where is the maximum intensity for the chaperone plus substrate mixture divided by the maximum intensity of the substrate alone. Values of 1 and 0 respectively indicate maximum protection and the absence of protein against aggregation.

**Supplementary Figure 2.**
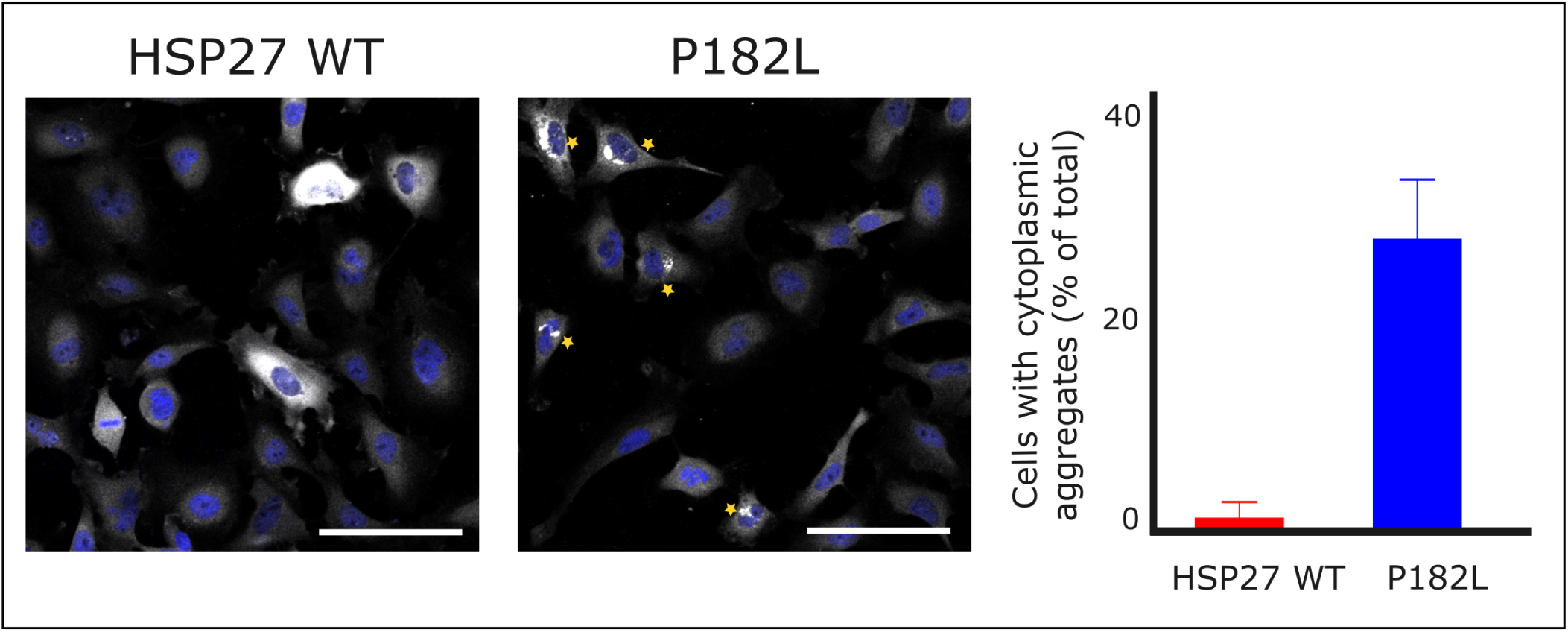
The P182L mutation increases the number of cells with large cytoplasmic insoluble aggregates upon HSP27 expression. Expression of V5-epitope-tagged HSP27 (WT or P182L mutant) in HSP27 knock-out HeLa cells and immunostaining of HSP27. All cells present in images of random large microscopic fields (640 µm × 640 µm) were visually scored for the presence of large high intensity aggregates. In total, 1361 cells were evaluated in 14 microscopic fields. The error bar represents the standard deviation between the different microscopic fields. Asterisks indicate cells with aggregates. Scale bar = 100µm.

**Supplementary Figure 3.**
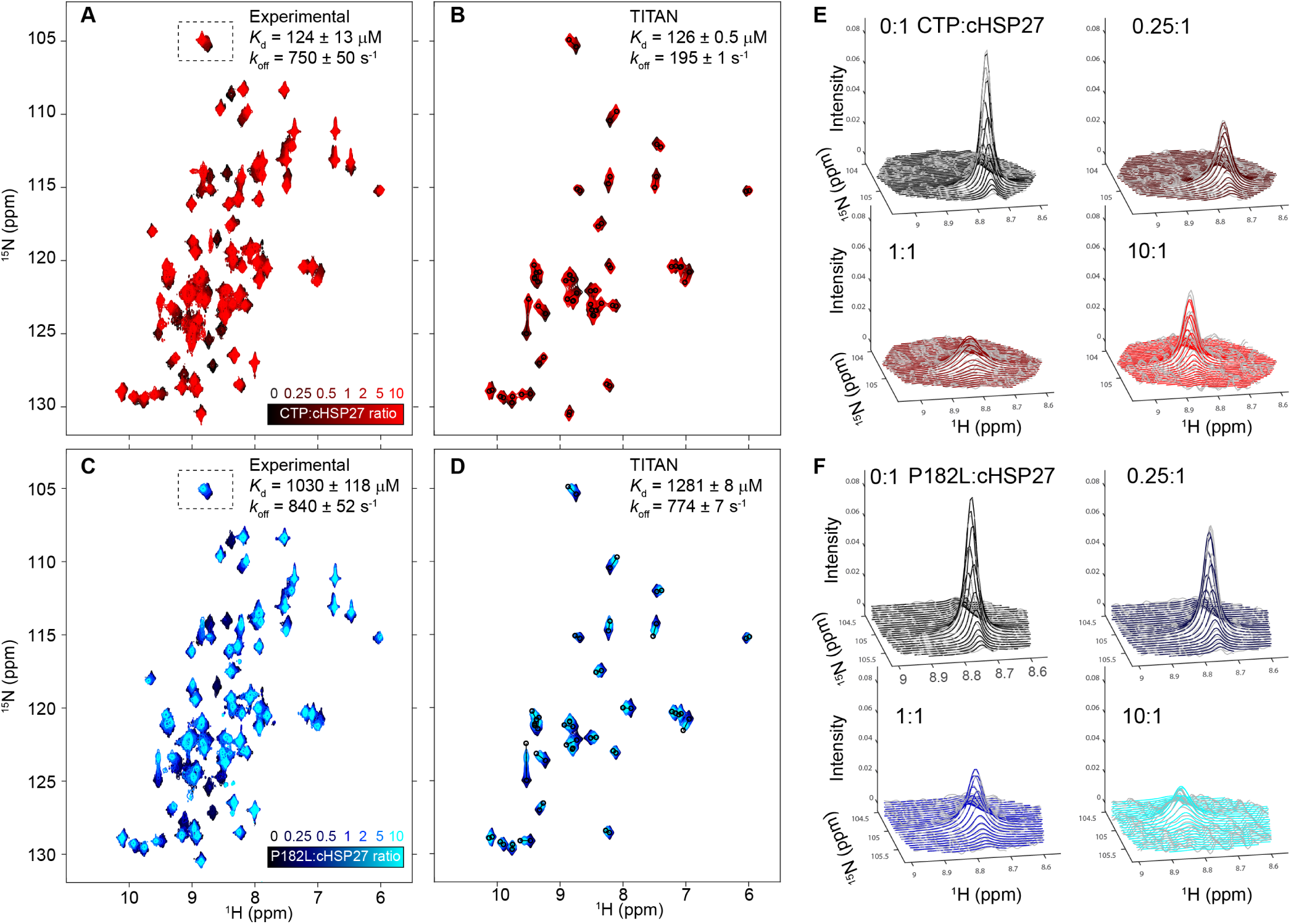
Analysis of the peptide binding NMR titration data with the TITAN software. (**A**), Experimental 2D ^1^H-^15^N HSQC spectra of ^15^N-ACD (black) in the presence of increasing amounts of unlabeled CTP (red color gradient). The globally determined *K*_*d*_ from fitting resonances in fast exchange to eqn 2 is listed in the upper-right corner. The *k*_*off*_ listed is derived from CPMG RD experiments (*vide infra*). (**B**) Using resonances in fast, intermediate, and slow exchange, the software TITAN fits NMR titration data to the Bloch-McConnell equations. For the selected resonances in the ACD, the reconstructed spectra are shown as fitted by the software. The *K*_*d*_ and *k*_*off*_ fitted by TITAN are listed in the upper-right corner. The color scheme is the same as panel A. (**C**) Same as panel A except for the peptide for the P182L variant. (**D**) Same as panel B for the peptide P182L. (**E, F**) For the resonances in panels A and C, selected peak shapes (grey lines) and the corresponding fits (colored lines) by TITAN are shown.

**Supplementary Figure 4.**
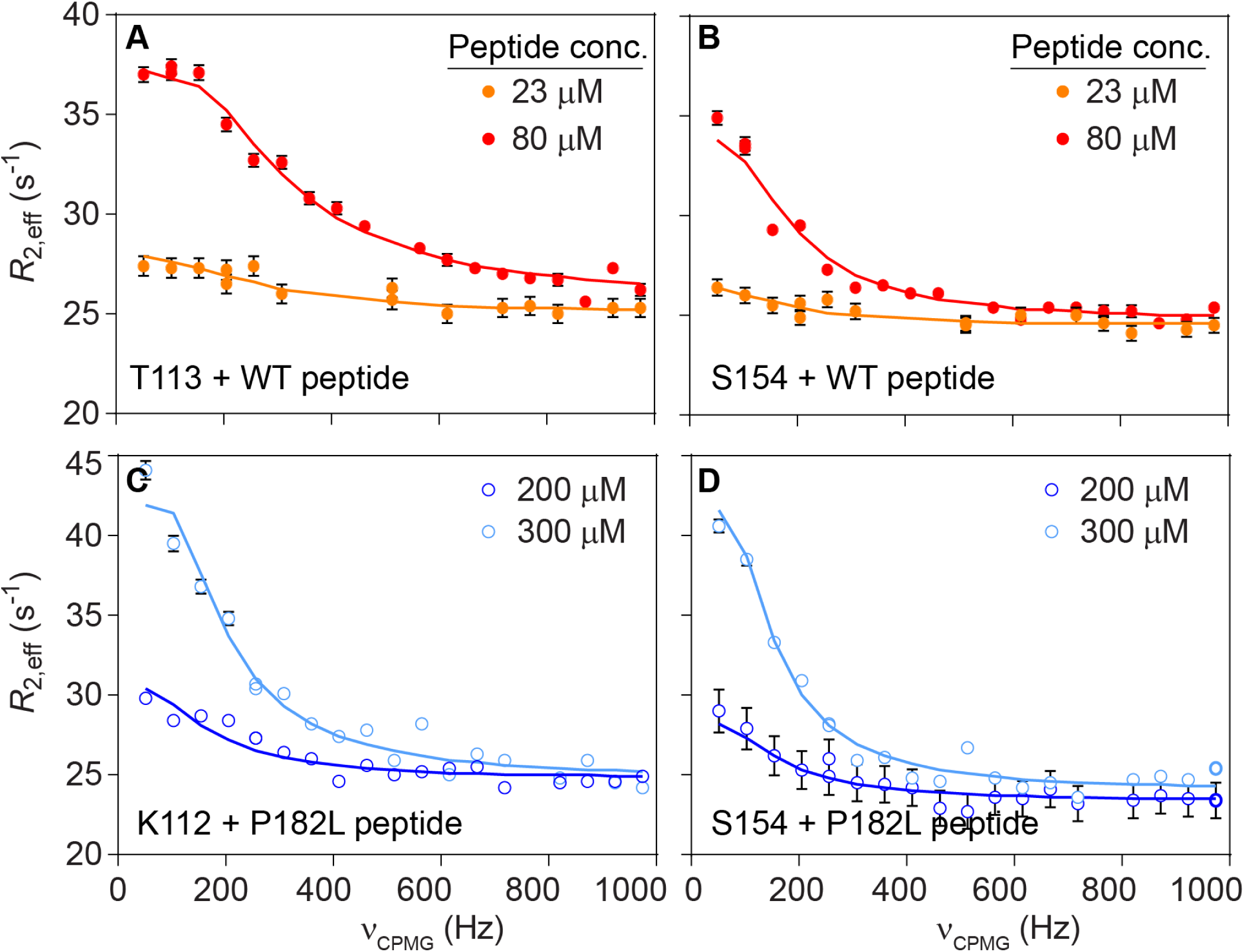
Concentration-dependence of dispersions from peptide-bound cHSP27. (**A, B**) i. 15N CPMG RD data for a residue in the β4 strand (T113) and the β8 strand (S154) in the presence of ii. ca. 2% (red) or ca. 0.2% cHSP27-WT peptide complex. (**C, D**) Dispersions from residues in the β4 (C) or β8 strand (D) as a function of added P182L peptide. In dark blue is ca. 2% of the cHSP27-P182L peptide complex and in light blue is ca. 4% of the cHSP27-P182L peptide complex.

**Supplementary Figure 5.**
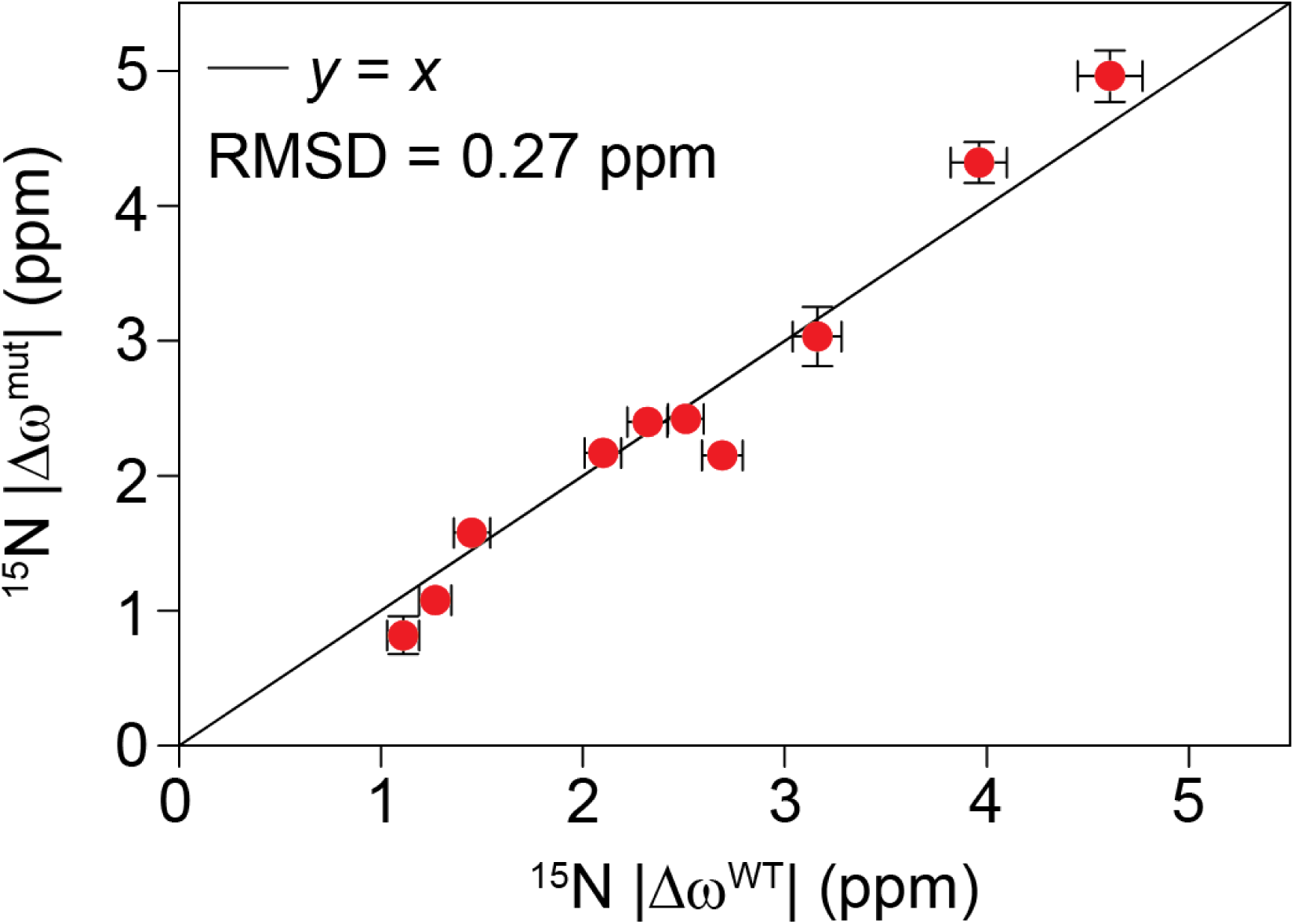
Similarity of the WT and mutant peptide-bound states. The overall similarity between the 15N |,Δω| values for the ACD in the presence of WT or P182L peptides (RMSD = 0.27 ppm), suggests that cHSP27 adopts a relatively similar conformation when bound to the WT or P182L peptide. The superscript mut refers to P182L. Note that most of these resonances were severely broadened in the peptide titrations in Fig. 3 of the main text, and thus their CSPs could not be directly measured. CPMG RD, however, enabled measurement of their 15N |Δω| values upon peptide binding.

**Supplementary Figure 6.**
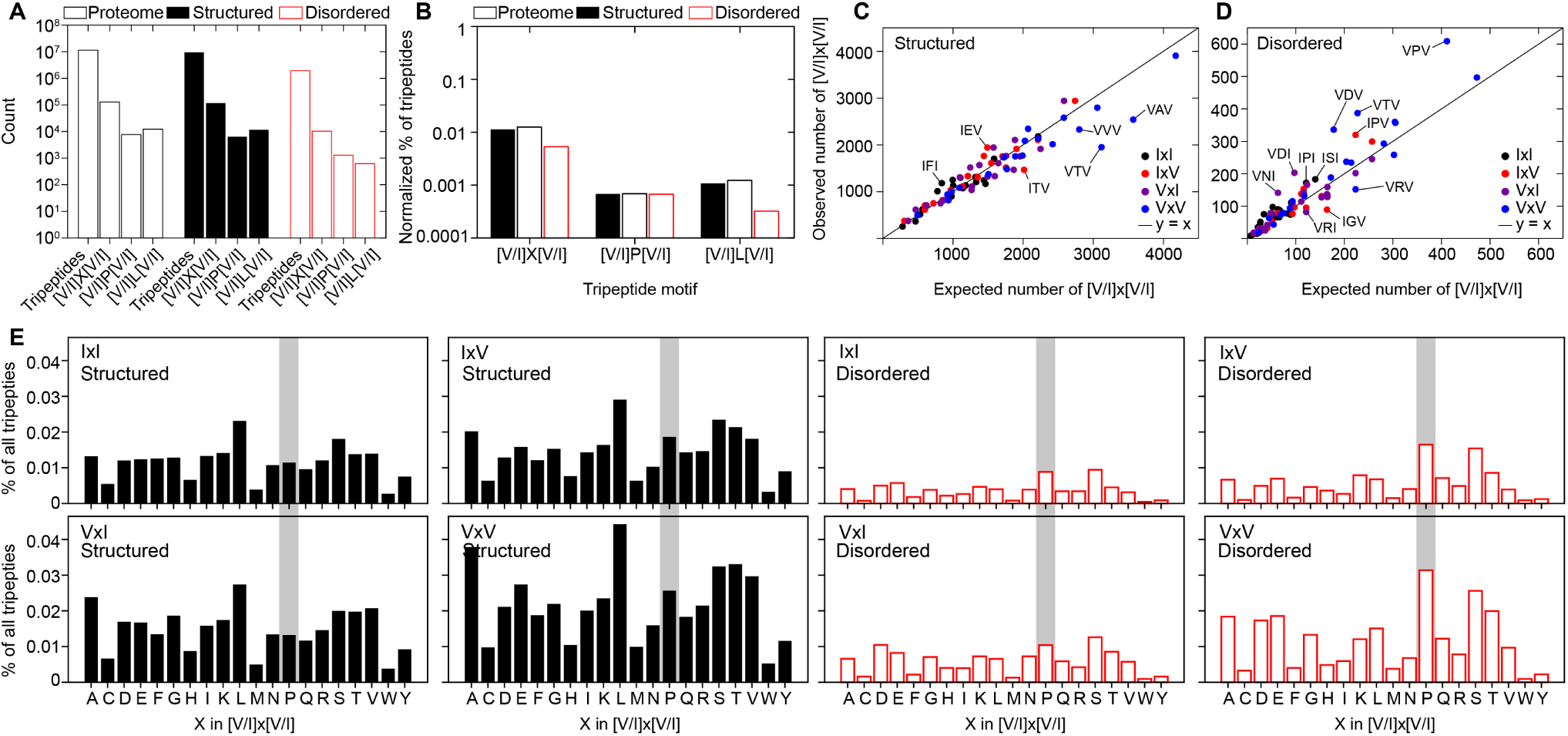
Bioinformatics analyses of [V/I]x[V/I] motifs in the proteome. (**A**) The total number of tripeptides and [V/I]x[V/I], [V/I]P[V/I], and [V/I]L[V/I] motifs in (empty black bars) the proteome, (black bars) structured regions of the proteome, and (empty red bars) disordered regions of the proteome. (**B**) The percentage of [V/I]x[V/I], [V/I]P[V/I], and [V/I]L[V/I] motifs for the proteome, structured regions, and disordered regions normalized to the total number of tripeptides present in each. (**C**) The observed number of IxI, IxV, VxI, and VxV motifs in structured regions of the proteome compared against the expected number of such motifs calculated using the frequency of amino acids. (**D**) The same as (C) but for disordered regions. Motifs that are either somewhat enriched or depleted are labeled. For the structured and disordered plots shown here, the *R*^2^ values from a linear regression (not plotted) are respectively 0.89 and 0.86 with slopes of 0.83 and 1.12. (**E**) The distribution of [V/I]x[V/I] motifs as a function of the central residue X for all of the possible combinations of [V/I]x[V/I]. Structured regions are shown in black bars and disordered regions in empty red bars. Proline is indicated with a grey box to denote the [V/I]P[V/I] motif.

**Supplementary Figure 7.**
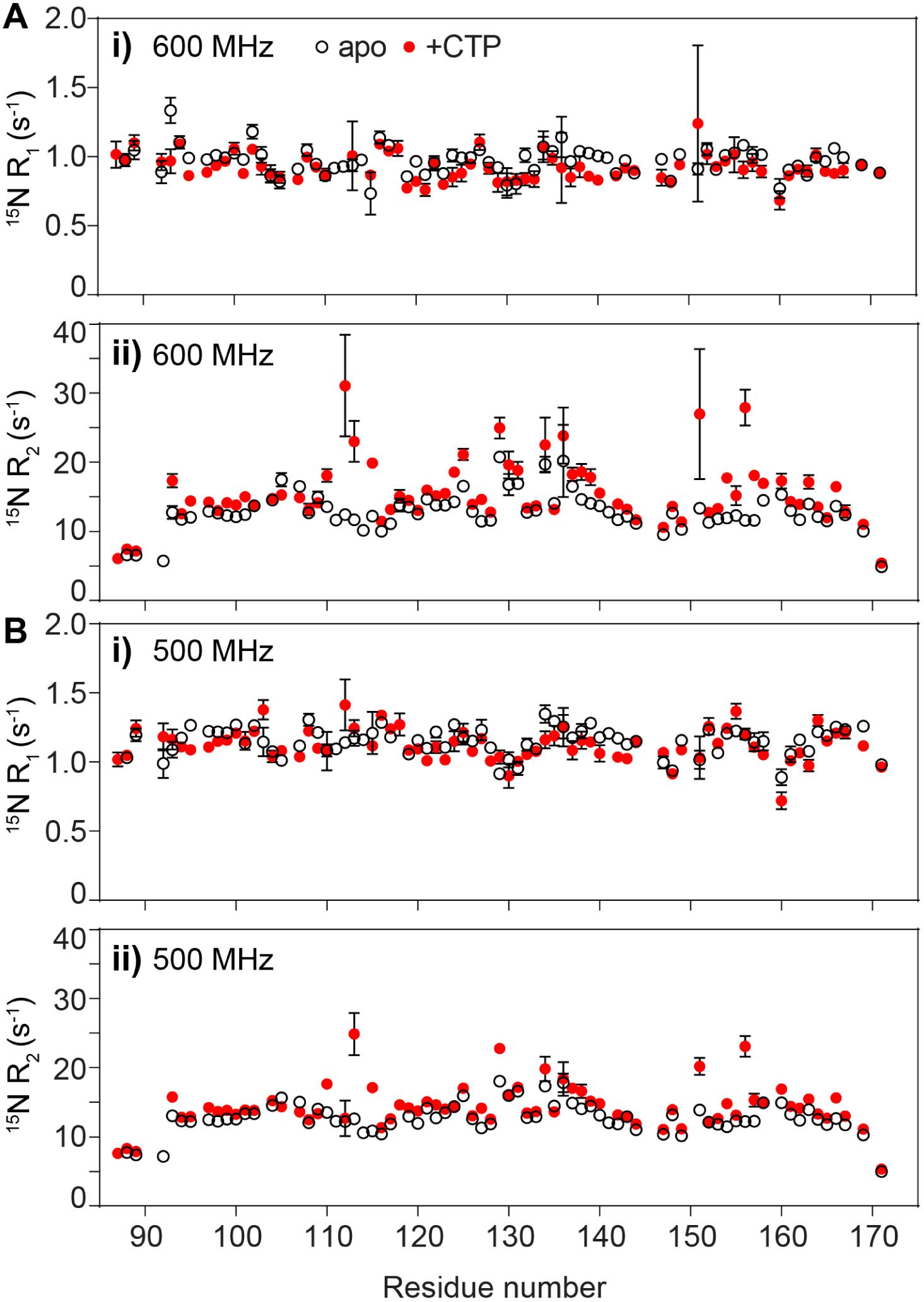
^15^N relaxation of peptide-free and peptide-bound cHSP27. ^2^H, ^15^N-labeled cHSP27 was prepared at a concentration of 0.3 mM in 30 mM sodium phosphate, 2 mM EDTA buffer at pH 7 and 37 °C. ^15^N *R*_1_ (i) and ^15^N *R*_2_ (ii) relaxation rates were measured at 600 MHz (A) and 500 MHz (B). In the figure, apo means peptide-free cHSP27 and +CTP refers to the peptide-bound form. To generate the peptide bound form, 3.8 mM WT peptide (CTP) was added to 0.3 mM cHSP27. For a *K*_*d*_ of 125 µM, this corresponds to 96.8% of bound cHSP27. The elevated *R*_2_ values in A i and B i reflect contributions due to chemical exchange between the free and bound forms.

## Notes

#### Summary of Updates

This version of the manuscript has been revised to update the following: small changes to the title and text; minor modifications to Figure 2, Figure 4, and Figure 5; and moving most of the Materials & Methods to the Supplementary Information.

